# DiPRO1 dependent transcriptional and epigenetic regulation distinctly controls the fate of muscle and mesenchymal cancer cells

**DOI:** 10.1101/2023.01.08.523169

**Authors:** Jeremy Rich, Melanie Bennaroch, Laura Notel, Polina Patalakh, Julien Alberola, Paule Opolon, Olivia Bawa, Windy Rondof, Antonin Marchais, Philippe Dessen, Guillaume Meurice, Melanie Porlot, Karine Ser-Le Roux, Nathalie Droin, Hana Raslova, Birgit Geoerger, Iryna Pirozhkova

**Affiliations:** UMR8126 CNRS, Gustave Roussy Cancer campus, Université Paris-Saclay, France; Pathology and Cytology Section, UMS AMMICA, CNRS, INSERM, Gustave Roussy Cancer campus, Université Paris-Saclay, France; Bioinformatics Platform, UMS AMMICA, CNRS, INSERM, Gustave Roussy Cancer campus, Université Paris-Saclay, France; Department of Pediatric and Adolescent Oncology of Gustave Roussy, U1015 INSERM; Pre-clinical Evaluation Unit (PFEP), INSERM, Gustave Roussy Cancer campus, Université Paris-Saclay, France; Genomics Unit, Department of Integrated Biology (PBI), Gustave Roussy Cancer campus, Université Paris-Saclay, France; UMR1287 INSERM, Gustave Roussy Cancer campus, Université Paris-Saclay, France; INSERM U1016, CNRS UMR 8104, Institut Cochin, Université Paris-Cité, France

## Abstract

We have recently identified the uncharacterized ZNF555 protein as a component of a productive complex, which is involved in the morbid function of the 4qA locus in facioscapulohumeral dystrophy. As a result of our current findings, ZNF555 is hereinafter referred to as DiPRO1 (Death, Differentiation and PROliferation related PROtein 1). In this study, we provide substantial evidence that DiPRO1 plays a role in human myoblast differentiation. It acts on regulatory binding regions of SIX1, which is a master regulator of myogenesis. We further describe the relevance of DiPRO1 in mesenchymal tumors, such as rhabdomyosarcoma (RMS) and Ewing sarcoma. DiPRO1 plays a repressor role in these tumors via the epigenetic regulators TIF1B and UHRF1 in order to maintain methylation of regulatory cis-elements and promoters. Loss of DiPRO1 eradicates cancer cells, by switching on a distinct transcriptional and epigenetic program. It consists of mimicking the host defense against the virus response by awakening the retrotransposable repeats (RE) and the ZNP/KZFP gene family. DiPRO1 also contributes to the balance of cellular decisions toward inflammation and/or apoptosis by controlling TNF-α via NF-kappaB signaling. Finally, we demonstrate that mesenchymal cancer tumors are vulnerable in response to si/shDiPRO1-based nanomedicines, positioning DiPRO1 as a potential new target for therapeutic intervention.

**Summary:** 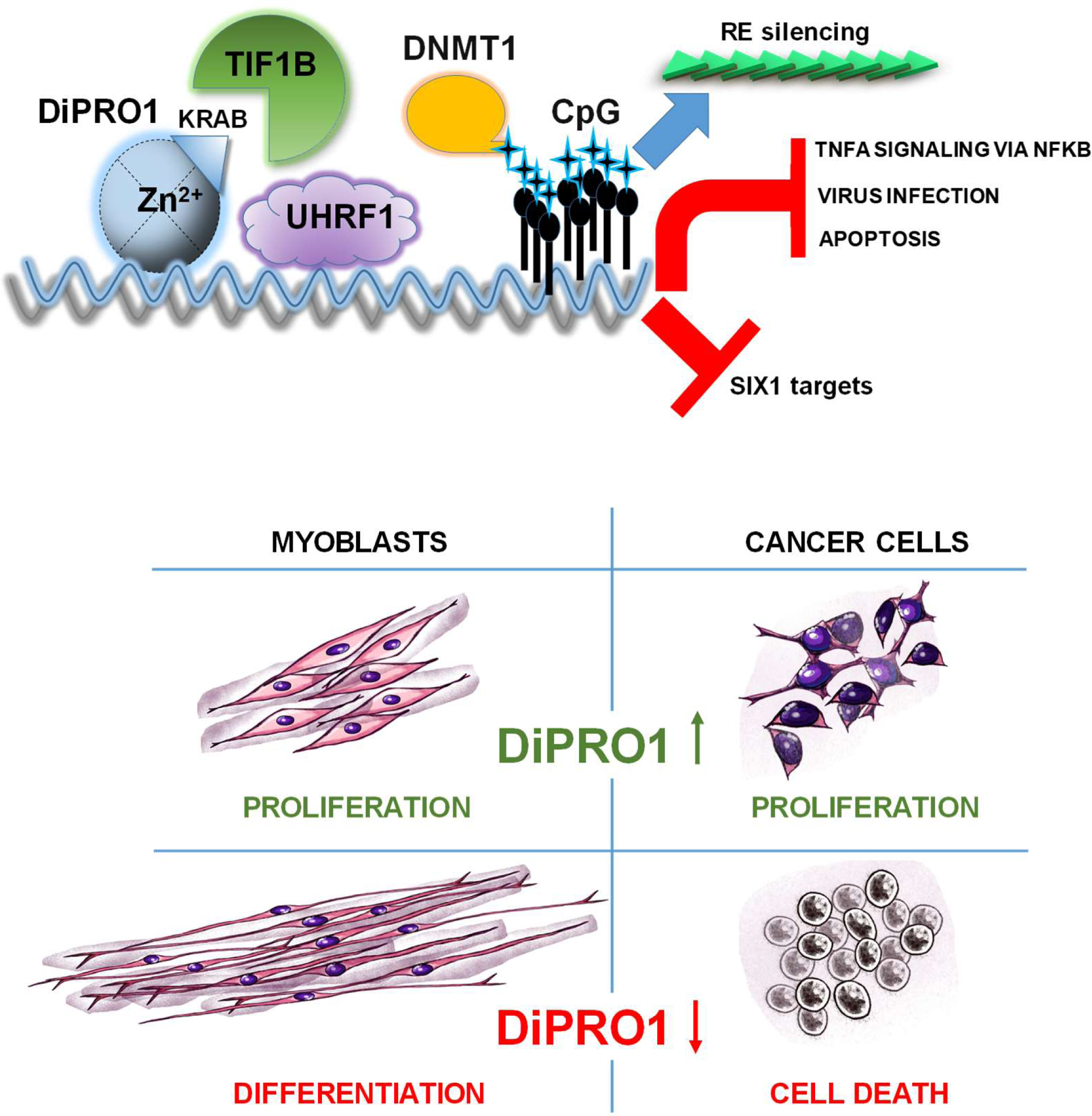

## INTRODUCTION

Scientific discoveries keep highlighting and clarifying the mechanisms controlling cell proliferation and differentiation. While the four transcription factors of the Yamanaka cocktail have been shown to be sufficient to induce pluripotency (1), further research has focused on the mechanisms of stem cell differentiation, their plasticity and trans-differentiation potential (2). Investigations of the reprogramming of cancer cells, which possess characteristics similar to those of stem cells, is a main topic in cancer research (3). They focus on factors that can control the trajectory of differentiation and induce tumor cells to behave less aggressively.

The zinc finger (ZNF) family is composed of 1723 annotated genes in the human genome (4). The zinc-finger domain is one of the most abundant DNA-binding motifs of eukaryotic transcriptional factors. The amino acid identity of the binding site defines the DNA targeting sequences of zinc fingers and their corresponding functions. Through their ability to regulate gene expression, ZNF proteins are involved in cell proliferation, differentiation and cancer progression (4, 5). The C(2)H(2) zinc fingers (6), CCCH zinc fingers (7) and the RanBP2 type zinc fingers (8) may be involved in RNA binding promoting regulation of mRNA expression or processing. Depending on the presence of protein-binding domains, such as BTB/POZ, Krüppel-associated box (KRAB) and SCAN, ZNF may be involved in protein-protein interaction (5). At least one-third of mammalian ZNF proteins include a KRAB domain. The family of KRAB-containing ZNF proteins (KZFP) is specific to tetrapod vertebrates and has been expanded extensively to hundreds of members along all mammalian evolution (9, 10). Many of the KZFP act in association with KRAB-associated protein 1 (protein: TIF1B/KAP1; gene: TRIM28) to repress transcription by recruiting histone deacetylases, SETDB1 histone methyltransferase, HP1, and the NuRD (nucleosome remodeling deacetylase) complex containing histone deacetylases, and may contribute to methylation of CpG islands (11, 12). One of the consequences of the KZFP repressive function is a silencing of repetitive DNA sequences, which constitute a large proportion of cis-regulatory elements, including enhancers, promoters, suppressors, insulators and TF binding sites (13, 14). Epigenetic fine-tuning of repetitive DNA sequences is essential for appropriate development and cellular homeostasis, whereas any deregulation can lead to aberrant dedifferentiation and various pathological processes, including carcinogenesis (15). Nevertheless, recent studies suggest that KZFP can associate with a variety of cofactors and play a dual role in transcription, extending the one previously believed to be limited to transcriptional repression (12, 16). Despite the high abundance of KZFP in the human proteome, the biological roles played by most of them remain unknown so far. Thus, the experimental demonstration of predicted ZNF gene products, as well as the study of their functions is an important area of biomedical research.

Recently, we identified the ZNF555 (Q8NEP9) protein as part of the productive transcriptional hub in myoblasts from patients with facioscapulohumeral muscular dystrophy (FSHD). This protein binds to a cis-regulatory element of the beta-satellite repeat (BSR) that controls transcription of muscle specific and pro-apoptotic gene ANT1/SLC25A4 (17). Related to the above findings, the present study reports novel myogenic functions for ZNF555. We demonstrate that ZNF555 shares regulatory binding sites with SIX1, an upstream regulator of myogenesis (18) and epithelial mesenchymal transition (19). ZNF555 drives a muscle differentiation inhibition through SIX1 targets. We therefore applied ZNF555 targeting first to rhabdomyosarcoma (RMS) and then to Ewing sarcoma cells, two mesenchymal cancers, which showed increased levels of ZNF555 expression. Excitingly, loss of ZNF555 results in cancer cell death. This was associated with global demethylation of CpG islands within the genes bodies and retrotransposable repeat elements (RE). Loss of methylation, in turn, may contribute to increased expression of pro-apoptotic genes as well as genes involved in inflammation and defense against viral infection.

The cell death effect was further validated by targeting ZNF555 in the human cell-derived xenograft (CDX) mouse cancer model. Finally, the pan-cancer analysis of primary pediatric tumors illustrates the potential clinical relevance of ZNF555 targeting for mesenchymal cancers and SOX2-related cancers. On the basis of morphological changes and transcriptional signature, ZNF555 was designated as Death, Differentiation and PROliferation-related PROtein 1 (DiPRO1).

## RESULTS

### DiPRO1 is an evolutionarily conserved gene in placental mammals

The DiPRO1/ZNF555 protein contains 628 amino acids (73.1 kDa, pI 8.9). It is encoded by NM_152791 and BC022022 [N107D H194N K515T] (Mammalian Gene Collection) with some amino acid variations. The DiPRO1 protein includes one KRAB A-box and fifteen C2H2 zinc finger domains. *In silico* prediction describes three isoforms: Q8NEP9 (628 AA), Q8NEP9-2 (543 AA) and Q8NEP9-3 (627 AA) produced by alternative splicing (Supplementary Fig. S1A). Isoform 2 lacks three zinc finger domains compared with isoforms 1 and 3. Isoform 3 has 99.8% similarity to isoform 1 with one missed amino acid difference. Protein blast analysis identified related proteins (BlastP, threshold 0.001) in vertebrates, with maximum homology in primates (98.7% in great apes and 89.5% in NW monkeys). However, a low homology of 50% does not allow identification of the DiPRO1 homologous protein in *Mus musculus*. Analysis of the phylogenetic history of all ancestral species showed the conservation of the DiPRO1 gene within placental mammals (Fig. 1A), indicating that DiPRO1 is an evolutionarily newly emerged gene found only in higher organisms (20). Significant (p<0.01) expansion of the DiPRO1 gene was observed in pigs, American black bears, and ferrets. During evolution, these species acquired two DiPRO1 genes. Consistent with the results of the protein blast analysis, the DiPRO1 gene is absent in rodents and rabbits, and also with the exception of the naked mole rat, indicating that these animal models are not suitable for the DiPRO1 study.

**Figure 1.**
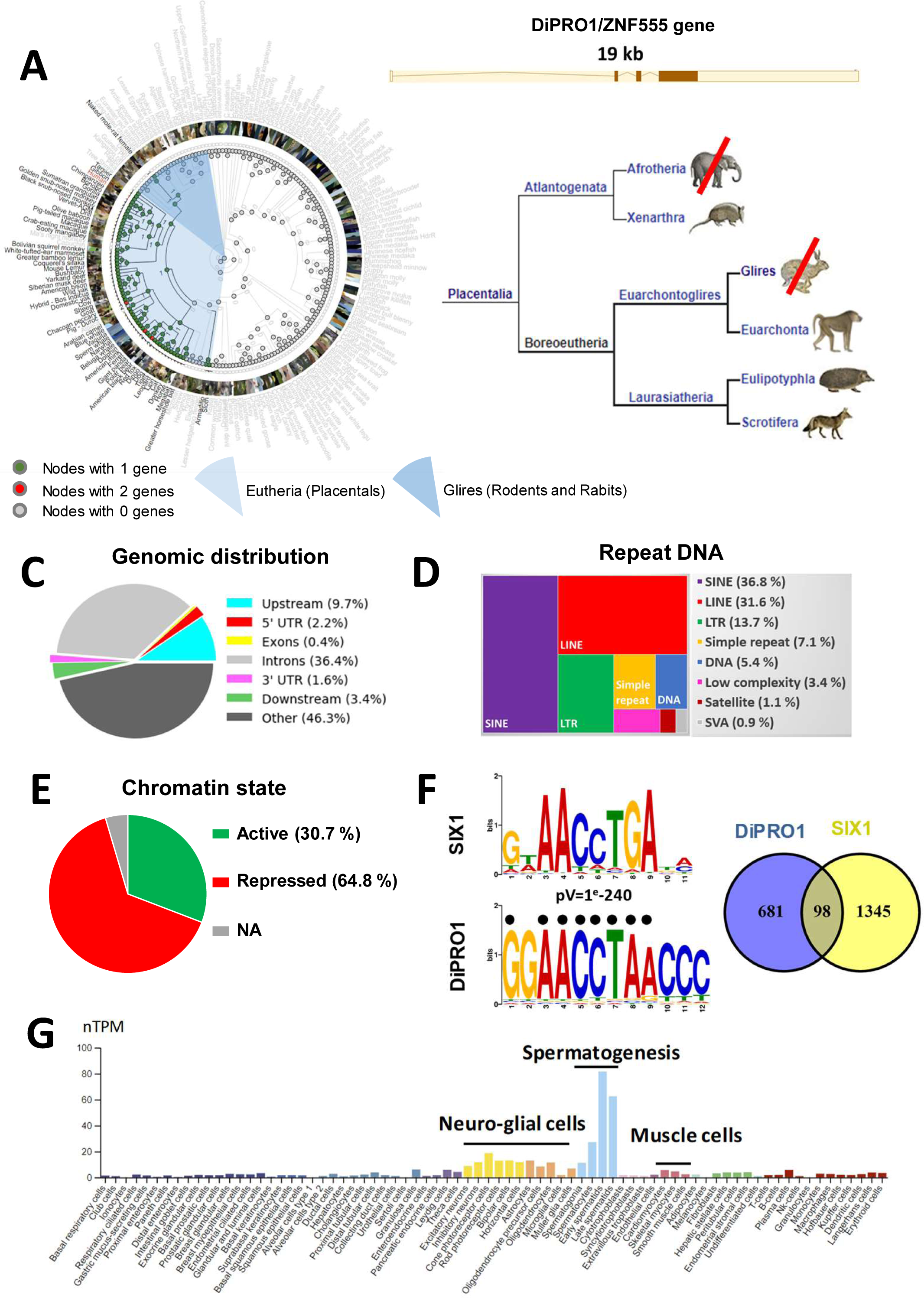
DiPRO1 direct targets, *de novo* motifs and associated chromatin state. **A,** Phylogenetic tree of an Ensembl gene family representing the evolutionary history of gain/loss events of the DiPRO1/ZNF555 gene (chr19:2.841.435-2.860.471 region of GRCh38). A branch was marked as an expansion or contraction if the p-value determined by the CAFE tool is <0.01. The numbers at each node refer to the number of DiPRO1 genes in the ancestral species. The color of geach node reflects the number of genes colored according to the legend below. Species names are colored; red (Homo sapiens for the ZNF555 gene), black (species with the ZNF555 gene in Ensembl in this tree), and gray (species with no ZNF555 gene in this tree). **C-F,** ChIP-seq data for DiPRO1 (GSM2466593) performed in HEK 293T cells. **C,** Pie chart of the distribution of DiPRO1 ChIP peaks relative to the gene body regions. The graph shows the percentage for each genomic location category. The upstream region was defined at -10Kb from TSS and the downstream region at 5Kb from TES. Distribution of DNA repeats (**D**) and chromatin status (**E**) in DiPRO1 binding regions **F,** The most specific DiPRO1 consensus motif 1 and its best match SIX1 (top left). The motif alignment was shown for the SIX1 human motif MA1118_Jaspar using the Tomtom tool. The black dots match the Six1 mouse motifs from Santolini et al. (DOI: 10.1093/nar/gkw512) and Liu et al (DOI: 10.1093/nar/gkq585). Gene overlap between the *cis*-regulatory regions of DiPRO1 and SIX1. **G,** Single cell type specificity of DiPRO1. Enhanced cell types are indicated. Data are taken from The Human Protein Atlas (www.proteinatlas.org).

### DiPRO1 binds the direct targets of SIX1

Recently, we have identified the DiPRO1 protein, as part of the productive transcriptional complex in the myoblasts of FSHD patients (17). In order to further explore the potential muscle-related functions and transcriptional co-regulators, we established the DiPRO1 direct physical targets by analyzing published ChIP-seq data (10). The genomic distribution of DiPRO1-associated regions showed that 54% of the interactions occurred within gene-related regions, from 10 kb upstream of the transcription start site (TSS) to 5 kb downstream of the transcription end site (TES) (Fig. 1C). A considerable fraction of all detected regions (69.5%) was observed in RE, including short dispersed nuclear elements (SINE), long dispersed nuclear elements (LINE), and long terminal repeats (LTR) (Fig. 1D). The chromatin state analysis based on histone marks (21, 22) in the binding regions of DiPRO1 demonstrated their association with both repressed (64.4%) and active chromatin (30.7%) states, including promoter, enhancer and transcribed regions (Fig. 1E, Supplementary Fig. S1B, Supplementary Data S1A). The Gene ontology (GO) analysis of direct transcriptional targets suggested that DiPRO1 functions may be coupled to cell cycle, transport, transcriptional co-regulator activity, and differentiation (Supplementary Fig. S1C). Among them, MYH2 and MYH3, classical genes of mature and embryonic skeletal muscle, FABP3 (23), NCOA2/TIF2 (24) and SIRT1 (25) are directly related to muscle functions. Interestingly, NCOA2 fusion with PAX3 has been shown to inhibit myogenic differentiation in muscle cancer (26).

*De novo* motif identification for DiPRO1 revealed the motif with the highest enrichment rate (motif 1, GGGTTAGGTTCC), which accounted for 16.0% of the target DiPRO1 sequences (Fig. 1F, Supplementary Fig. S1D). The best match to this motif corresponds to the transcription factor SIX1 (27–29). In addition, the most abundant motif (motif 8, CAGAGGCGCACA) was found in 37.8% of the DiPRO1 target sequences, showing the highest homology to the ZSCAN4/GM397 motif (Supplementary Fig. S1E). The most proximal genes associated with the *cis*-regulatory regions of DIPRO1 binding were shared between those of SIX1 (12.6%, jaspar.genereg.net) and ZSCAN4 (77.7%, GSE105311) (Fig. 1F, Supplementary Data S1B), thereby supporting the motif discovery analysis. Remarkably, the SIX1 homeoprotein is an upstream regulator of myogenesis and a repressor of RMS muscle cancer differentiation (18,30–32).

The functions of ZSCAN4 are not specifically related to muscle, but rather to the epigenetic regulation of pluripotency in normal and cancer stem cells, affecting their growth (33, 34). Interestingly, ZSCAN4 has been found to be associated with microsatellite DNA regions (35) and upregulated in muscles expressing a long FSHD-related macrosatellite transcript (DUX4-FL) (36). In relation to RE, we have previously demonstrated that DiPRO1 can physically interact with a β-satellite DNA repeat (4qAe) in FSHD muscle cells (17). We show that the DiPRO1 binding motif 8, homologous to ZSCAN4, matches the repeat sequence. Consistently, the motif prediction was in good agreement with the experimental data (Supplementary Fig. S1F). To further investigate transcriptional co-regulation of DiPRO1, the ENCODE collection of ChIP-seq data was used (37, 38). The analysis revealed several regions of joint binding of DiPRO1 and TIF1B/KAP1, a mediator of transcriptional repression by KZFP (Supplementary Data S1A) (12, 39).

Collectively, this analysis predicts that DiPRO1 is a transcriptional regulator acting through gene promoters and *cis*-regulatory RE. DiPRO1 shows overlapping binding with SIX1, further arguing that it may play a role in myogenesis.

### DiPRO1 is a regulator of myogenic program in human myoblasts

Previously, we have shown an robust expression of DiPRO1 in myoblast cells in comparison to fibroblasts and its relevance to the muscular dystrophy FSHD (17). We have also reported that the most significant DiPRO1 expression fluctuations are related to musculoskeletal and connective tissue diseases according to the Disease Atlas (nextbio.com). We then queried a single cell RNA-seq database (www.proteinatlas.org) to assess DiPRO1 expression in normal cells. The DiPRO1 expression is enhanced in three major cell types, namely, germ cells, neuro-glial cells and muscle cells (Fig. 1G), supporting the prediction of DiPRO1 role in muscle. Therefore, we studied the role of DiPRO1 in muscle cells.

First, we performed DiPRO1 overexpression experiments. The full-length ORF (1887 bp) of DiPRO1 was fused to the C-terminus of the triple-tagged Flag, 6xHis, HA (FHH) YFP protein (pDiPRO1) within the pOZ retroviral backbone (Supplementary Fig. S2A). In addition, the DiPRO1 ORF was codon-optimized, which enabled us to distinguish between endogenous and recombinant DiPRO1 expression (Supplementary Fig. S2B). Using stable transduction of the pDiPRO1 retroviral vector, immortalized human myoblasts overexpressing the DiPRO1 mRNA and protein were generated and used for further study (Fig. 2A-B). The pDiPRO1 myoblasts were maintained in the myoblast proliferation medium. They had a rounded or short spindle shape with no alignment upon aggregation, in contrast to the long spindle or polygonal aligned myogenic parental cells (40, 41) (Fig. 2C). May-Grünwald-Giemsa (MGG) staining revealed that these cells had reduced size, hyperchromatic nuclei, and increased nucleus-to-cytoplasm (N/C) ratio (Fig. 2D), which are characteristics of undifferentiated and immature cells (42, 43). In addition, the pDiPRO1 myoblasts showed a significant increase in proliferation rate, comparable to that of RMS cells (Fig. 2E), and PAX3/6/9 gene upregulation (Supplementary Fig. S2C-E). In the differentiation medium, we did not observe multinucleated and elongated myotubes in the myoblasts overexpressing DiPRO1. The cells retained their disorganized shape and did not express muscle tropomyosin protein, suggesting a block in differentiation and fusion (Fig. 2F). To compare the effect of up-regulation of DiPRO1 with that of down-regulation of DiPRO1, myoblasts were then stably transduced with a lentiviral vector expressing a shRNA targeting DiPRO1 (shDiPRO1) (Fig. 2G). The resulting depletion of DiPRO1 induced the onset of cell cycle suppression at G1/S, with no difference in mortality compared to control cells (Fig. 2H, Supplementary Fig. S2F). One week later, the DiPRO1-depleted myoblasts underwent transformation into elongated, thin cells (Fig. 2I), showing increased MYOD, MYH1, and MYOG mRNA levels (Fig. 2J).

**Figure 2.**
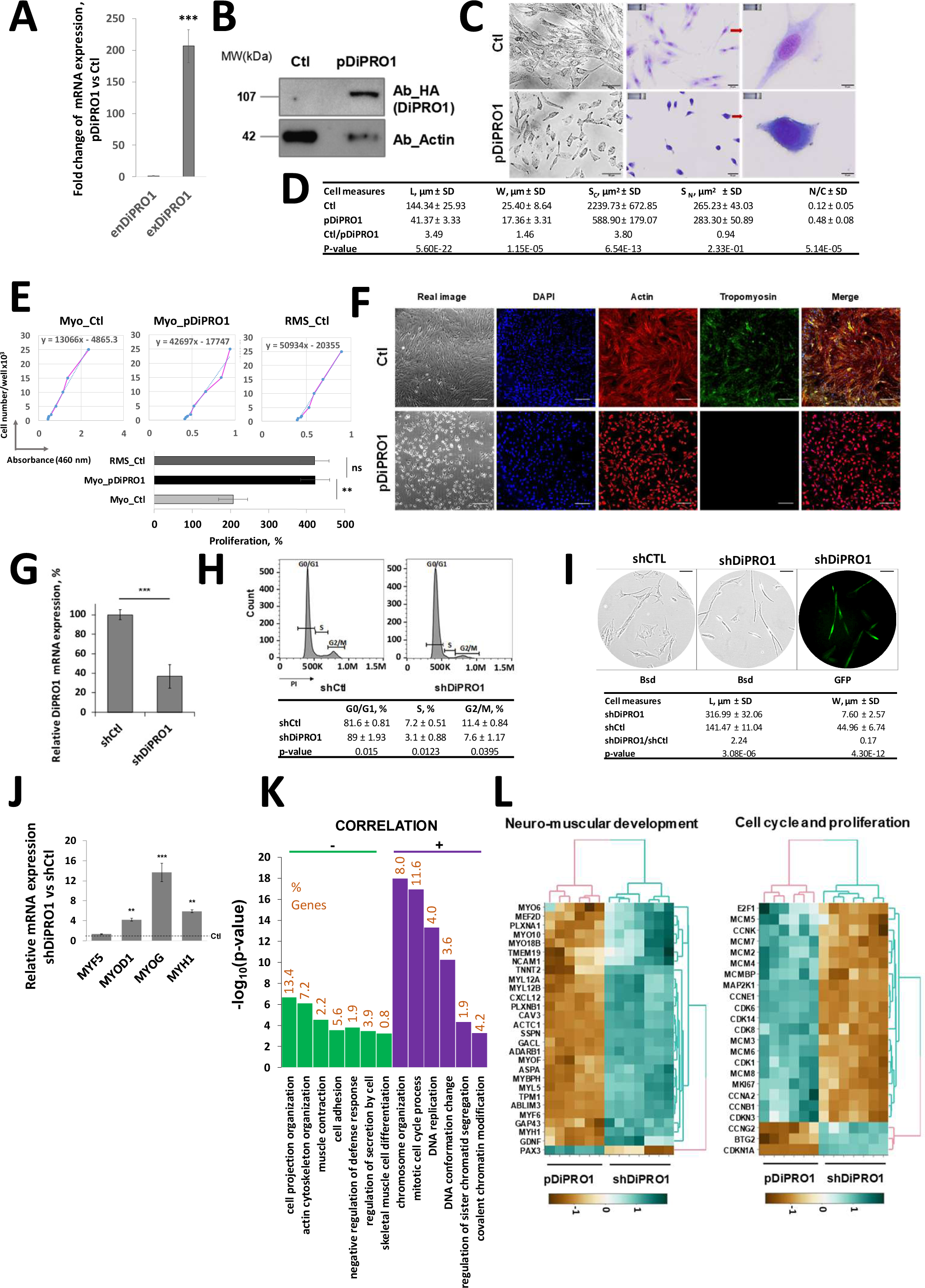
Genetic manipulation targeting DiPRO1 modulates the differentiation state of myoblasts. **A-G,** Two myoblast cell lines were transduced with a retroviral vector expressing the DiPRO1 ORF (pDiPRO1) and compared with parental control counterparts (Ctl). The efficiency of exogenous DiPRO1 overexpression was verified by mRNA expression analysis **(A)** and Western blotting **(B)**. The qRT-PCR results were normalized to the GAPDH gene and represent three independent total RNA extractions measured in triplicate, mean ±SD. Ct=45 was taken for exDiPRO1 ORF expression in control cells. The P-value indicates the difference between endogenous (enDiPRO1) and exogenous (enDiPRO1) DiPRO1 expression. Immunoblotting assays were performed with whole cell extracts. DiPRO1 was tagged with the HA-tag. Actin was served as an internal control. **C,** Morphological changes in myoblasts stably expressing pDiPRO1. Real images (left panel) and cells stained in MGG reagent (middle and right panels) are shown. Scale bar: 50 µm (left, middle) and 10 µm (right). **D,** DiPRO1 overexpression induces changes in cell dimension. L: cell length, W: cell width, Sc: cytoplasmic surface, Sn: nuclear surface, N/C: nuclear-cytoplasmic ratio. Unit of length = µm. Results represent m ± SD, n=20. **E,** DiPRO1 induces myoblast proliferation. The proliferation rate was estimated for DiPRO1-overexpressing myoblasts (Myo_pDiPRO1) and their control (Myo_Ctl) and compared with RMS cells (RMS_Ctl) by Counting Kit 8 (Sigma-Aldrich). Cell number was determined using a titration curve performed at different cell dilutions from 0 to 25,000 cells/well (0, 1.0×10^4^, 1.5×10^4^ and 2.5×10^4^). Absorbance was measured at 460 nm using a plate reader one hour after plating. Viable cells after 72 hours of proliferation were compared to the initial number of cells. Data are expressed as mean % relative to initial cell number ± SD and represent three independent experiments. **F,** Overexpression of DiPRO1 results in the block of myoblast differentiation. Cells were stained with anti-tropomyosin (green), anti-actin (red) and DAPI (blue) antibodies. The merge contains the combined images of three different stainings. Scale bar 120 µm. **G-J,** Two human myoblast cell lines were transduced with a lentivirus vector expressing a shRNA targeting the DiPRO1 gene (shDiPRO1) and compared with myoblasts transduced with the equivalent vector expressing a nontargeting shRNA (shCTL). G, DiPRO1 mRNA inhibition was verified by RT-qPCR analysis. Data were normalized to GAPDH expression and DiPRO1 expression in control cells was set to 100%, m ± SD, n=6 corresponding to two independent experiments. **H,** Cell cycle analysis of DiPRO1-depleted myoblasts was performed one week after transduction. Transduced myoblasts were stained with propidium iodide (PI). The percentage of dead and viable cells in each phase, according to DNA content, was determined by flow cytometry and compared with control cells, m ± SD, n=3 corresponding to three independent experiments. I, Morphological changes in myoblasts resulting from DiPRO1 knockdown (shDiPRO1) one week after transduction (top). Transduction efficiency was verified by confocal microscopy using the shDiPRO1 vector expressing GFP (top left). Scale bar 100 µm. Length (L) and width (W) of cells were measured (bottom). Length unit = µm. Results are presented as mean ± SD, number of fields n=20. (A-J) The t -test was applied for statistical analysis. *, **, *** indicate significant difference from the corresponding control, P < 0.05, 0.01, 0.001, respectively. **J,** DiPRO1 KD contributes to myogenic gene expression. Results of RT-qPCR analysis were normalized to GAPDH expression and presented as ratios to shCtl. The t -test was applied for statistical difference with the appropriate control, m ± SD, n=6 corresponding to two independent experiments. **K-L,** Global transcriptome analysis by microarray was implemented using total mRNA extracted from eight myoblast cell lines: with DiPRO1 knockdown (shDiPRO1) or overexpression (pDiPRO1) and corresponding controls. Three or four separate extractions were performed to collect replicates (n = 6-7 per condition). Differentially expressed genes in pDiPRO1- and shDiPRO1-myoblasts compared with corresponding controls showing negative and positive correlation with DiPRO1 expression (P < 0.05). The Limma R package was used for statistical analysis. **K,** GO terms of significantly regulated biological processes are summarized using the genetic interaction network. P < 0.01, kappa score threshold 0.4. **L,** Gene signatures of neuromuscular development, and cell cycle and proliferation pathways correlated with DiPRO1 expression versus control.

Next, genomic transcriptomic analysis was performed to identify differentially expressed genes (DEGs) focusing on common responders in the DiPRO1 overexpression and knockdown models (Supplementary Fig. S2C-E, G-H). Negatively correlated genes with DiPRO1 expression in loss-of-function and gain-of-function have been involved in muscle tissue development, actin cytoskeleton organization, and muscle contraction (Fig. 2K). Among them, MYH1, MYF6, MYO18B, TNNT2, and TPM1, key genes for skeletal muscle fiber development and muscle differentiation (41, 42) were identified (Fig. 2L, Supplementary Fig. S2I), which showed inversed expression with DiPRO1 gene levels. In contrast, PAX3, a marker of myogenic progenitor cells (43), showed a parallel correlation with DiPRO1 expression. The genes positively correlated with the DiPRO1 expression level were enriched in the cell cycle and replication pathways. The potential function of DiPRO1 in proliferation and cell cycle was reflected by transcriptional changes in genes encoding MCM and Ki-67 proteins, conventional markers of cell proliferation (Fig. 2L) (44, 45). The expression level of cell cycle regulatory genes was mainly distinguished by a positive correlation with cyclins A, B, E, cyclin-dependent kinases (CDK4, CDK6, CDK1), E2F1, and by a negative correlation with p21/CDKN1A, predominantly interfering in the G1/S phase transition, in agreement with the cell cycle analysis results (Fig. 2H). Interestingly, the global transcriptome of shDiPRO1 myoblasts differed less from controls (PC2=10.1%) than that of pDiPRO1 (PC1=34.1%), suggesting less severe rearrangement during DiPRO1 downregulation (Supplementary Fig. S2J). Taken as a whole, the functional models provide evidence that DiPRO1 maintains myogenic stem cells in a proliferative program and antagonizes their differentiation.

### DiPRO1 depletion induces mesenchymal cancer cell death *in vitro*

The RNA-seq analysis of 1375 human cancer cell lines (DepMap portal, Broad Institute) demonstrated particularly high expression of DiPRO1 in RMS (muscle cancer) and Ewing sarcoma (bone or soft tissue cancer), both of mesenchymal origin (Supplementary Fig. S3A) (46, 47). It has been previously reported that mesenchymal stem cells (MSCs) exhibit a high potential for transformation into mesenchymal tumors (48) and could be a possible source of trans-reprogramming of RMS into Ewing sarcoma by introducing the EWS-FLI1 gene (49). We therefore investigated whether the phenomenon of cell cycle restriction and proliferation inhibition induced by DiPRO1 KD in myoblasts could be extrapolated to mesenchymal cancer cells. Initially, TE671 RMS cells were subjected to DiPRO1 KD by transduction with lentiviral vectors each expressing one of the three shRNAs (shDiPRO1-1/2/3) or a non-targeted shCtl (Fig. 3A). Propidium iodide (PI) staining showed a 24% increase in the TE671-sub-G1 fraction compared with control 48 hours after transduction, indicating the onset of cell death (Fig. 3B).

**Figure 3.**
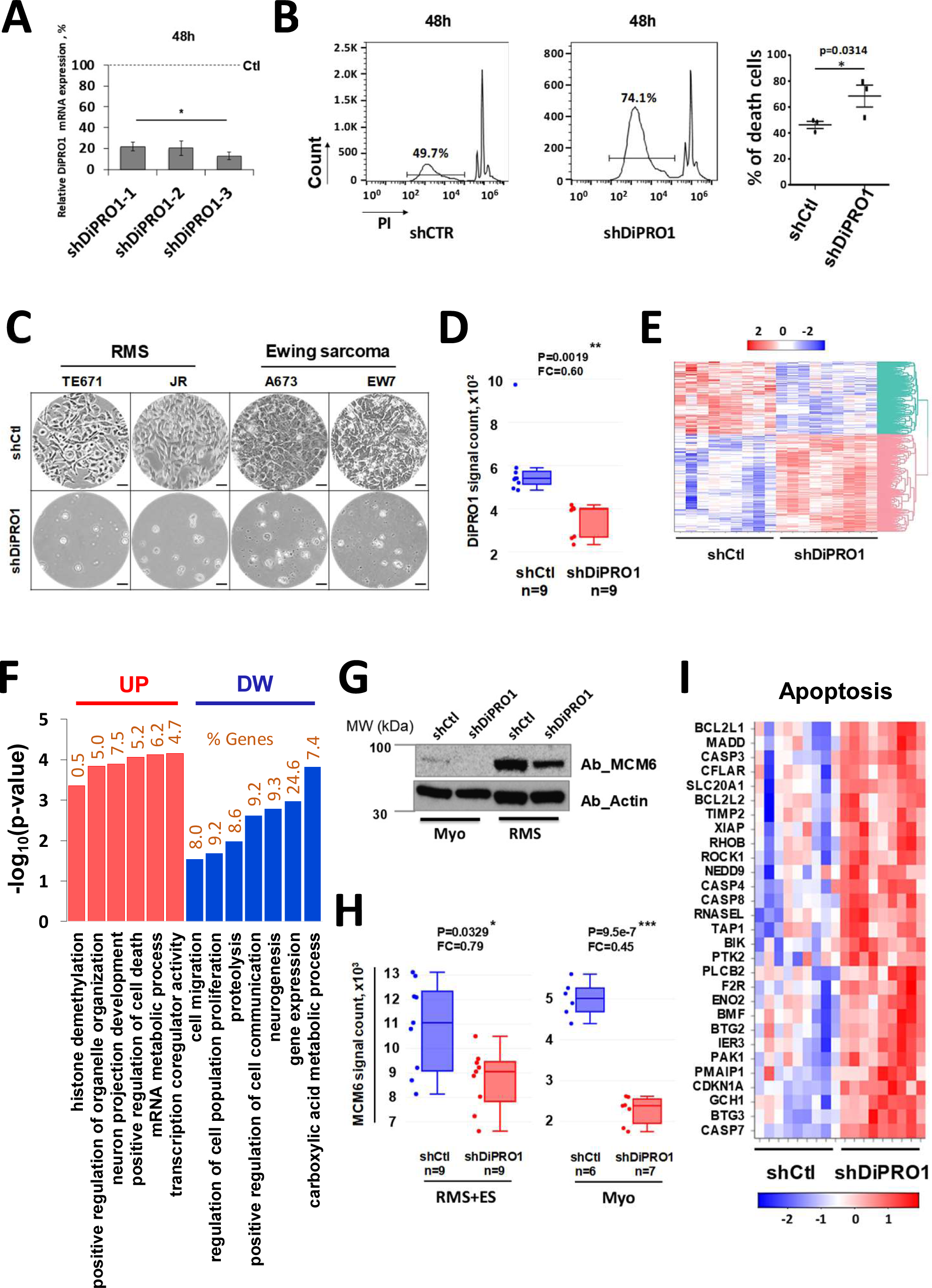
Functional depletion of DiPRO1 gene expression compromises cell fate and modulates the transcriptome of RMS and Ewing sarcoma cells. DiPRO1 knockdown was achieved by lentiviral transduction of vectors producing a non-targeted shRNA (shCtl) and a shRNA targeting the DiPRO1 gene (shDiPRO1). **A,** The efficiency of DiPRO1 inhibition was verified by mRNA expression analysis in TE671 cells. Three different shRNAs (shDiPRO1-1/2/3) were tested individually and the inhibition effect was analyzed by RT-qPCR. Total mRNA was extracted 48 h after transduction. The results represent at least three independent assays and were compared to shCtl, referenced as 100%, using the t-test. **B,** Induction of cell death in TE671 cells lacking DiPRO1 48h after transduction. Three independent experiments correspond to three individual shDiPRO1. Transduced RMS cells were stained with propidium iodide (PI). The percentage of dead cells in non-gated areas was determined by flow cytometry. The pairwise t -test was applied for statistical analysis. **C,** Dramatic cell death was observed 5-7 days after transduction of TE671 and JR RMS, and A673 and EW7 Ewing sarcoma cell lines. Scale bar: 50 µm. **D,** Boxplots of DiPRO1 downregulation in both RMS and Ewing sarcoma cells with DiPRO1 knockdown (red) and in control (blue). Data are presented as median ± dispersion, n=9. **E,** Heatmap showing genes that were affected in RMS and Ewing sarcoma cells under DiPRO1 knockdown (n=9) versus control (n=9), Padj<0.05. Clustering analysis of Euclidean distribution, full linkage. **F,** GO analysis of significantly downregulated (DW) or upregulated (UP) common DEGs in RMS and Ewing sarcoma cells compared with corresponding controls. FDR < 0.05, kappa-score threshold 0.4. **G-H,** MCM6 gene (**G**) and protein (**H**) expression in myoblasts (Myo) and RMS cells with DiPRO1 KD and control counterparts was determined 48 h after transduction. The mRNA expression data are presented as median ± dispersion. Western blot was performed using anti-MCM6 and anti-actin antibodies (internal control). **I**, Transcriptome of common apoptosis hallmarks in RMS and Ewing sarcoma cells. **D-F, H-I:** Global transcriptome analysis by microarray was implemented using total mRNA extracted from RMS and Ewing sarcoma cell lines: with DiPRO1 knockdown (shDiPRO1) and corresponding controls. Three to six separate extractions from two independent experiments were performed to collect replicates (n = 9 per condition). Differentially expressed genes in shDiPRO1-cells compared to corresponding controls (P < 0.05) were analyzed using the Limma R package.

This led to a progressive deregulation of the cell cycle as seen in the diploid phases G0/G1 and S (Supplementary Fig. S3B) (50), resulting in 95-100% cell death by day 5-7 (Fig. 3C). The observed effect was similar for three individually transduced shDiPRO1 (Supplementary Fig. S3C). RMS in children occurs as two main subtypes, embryonal (ERMS) and PAX-FOXO fusion positive alveolar (ARMS) (89). The TE671 cell line belongs to the ERMS subtype. Therefore, we additionally screened the ARMS JR cell line as well as the Ewing sarcoma cell lines (A673, EW7). As shown in Figure 3C and Supplementary Fig. S3D-F, the cell death phenomenon in sarcoma cells was comparable to that observed in the ERMS cells and was independent of the sarcoma type. Given the reproducibility of the effect induced by each shDiPRO1, one of the three shDiPRO1s (shDiPRO1-1) was used in subsequent experiments. It should be noted that the effect of DiPRO1 KD differed from that observed in myoblasts, suggesting the induction of an aberrant death program in malignant cells.

To identify shared anti-tumorigenic mechanisms of DiPRO1 depletion in mesenchymal cancer cells, we studied global gene expression changes in RMS TE671 and Ewing sarcoma A673 cells. The early molecular events (48h) of shDiPRO1-introduced cells were compared to those of shCtl. DiPRO1 depletion altered the expression of 4202 unique genes across these cells, corresponding to 57% of up- and 43% of downregulated genes (Limma, padj > 0.05) (Fig. 3D-E). This effect was coupled with the repression of genes involved in proliferation and apoptosis (Fig. 3F). One of the common responders to DiPRO1-downregulation in myoblasts and mesenchymal cancer cells was MCM6 (Fig. 3G-H), thus substantiating a common DiPRO1 role in cell proliferation.

Unlike in myoblasts, loss of DiPRO1 promoted upregulation of genes that are hallmarks of apoptosis (51), such as ROCK1 (52), RHOB (53), CDKN1A/p21 and BIK (54). Induction of the expression of the initiator caspase CASP8, the effector caspases CASP3 and CASP7, and the inflammatory caspases CASP4 and CASP5 suggests that depletion of DiPRO1 can induce caspase-dependent cell death (Fig. 3F, I) (55).

Collectively, these results demonstrate that DiPRO1 inhibition attenuates cell proliferation and induces apoptosis in mesenchymal cancer cells, highlighting an aberrant transcriptional program and associated functional effect in normal and malignant muscle cells.

### Proof of concept: *in vivo* antitumor activity evaluation

Tailored nanocarriers have gained huge research focus for tumor drug delivery (56). The branched and linear polyethylenimines (PEIs) have become prominent gene carriers for cancer and are explored in clinical trials (ClinicalTrials.gov NCT00595088, NCT02806687). To confirm our *in vitro* findings and to address the significance of DiPRO1 KD *in tumor* development in vivo, we evaluated the tumor growth when DiPRO1 was locally attenuated using sh/siRNA-based nanomedecines. Antitumor activity of the si- or shDiPRO1/jetPEI® at equimolar doses was investigated against Ewing sarcoma cell xenograft compared to scramble siRNA/jetPEI® (Ctl) in three independent experiments. The tumor cell internalization of siDiPRO1/jetPEI® coupled with Cy5 in living mice and extracted tumors was confirmed 24 and 72 hours after a single injection of 0.5 mg/kg (Fig. 4A, Supplementary Fig. S4A) and DiPRO1 gene inhibition was verified (Fig. 4B). In the first experiment (EW1), treatment was initiated when tumors had reached tumor volumes of 168±76 mm^3^ and the nanocomposites were administered according to a twice-weekly schedule at a dose of 0.5 mg/kg/injection during one week followed by 1 mg/kg/injection (Supplementary Fig. S4B). The treatment of both siDiPRO1 and shDiPRO1 nanomedicines showed moderate tumor growth stabilization as compared to controls in this rapidly growing model, however tumor regression and survival advantage was evident in smaller tumors (Supplementary Fig. S4C-E). Two additional independent experiments (EW2 and EW3) were performed starting treatment at tumor volumes of 81 ± 21 mm3 with an intensified three-times-per-week regimen during the first two weeks followed by twice-weekly 0.5 mg/kg/injection doses (Supplementary Fig. S4F). Visible ulcerations were observed in the EW3 model independent of treatment, therefore all experimental animals were sacrificed at day 12 according to animal protection endpoints and the survival analysis was not taken into account. In three independent experiments, the treatment with si- or shDiPRO1/jetPEI®-nanomedicines reduced tumor growth starting from day 4 of treatment (Fig. 4C and Supplementary Fig. S4C-D, G-H, J-K). Both treatments resulted in the survival advantage (p < 0.01, Mantel-Cox test) (Fig. 4D and Supplementary Fig. S4E,I). Tumor growth delay (TGD) and tumor growth inhibition at day 10 (TGId10) was more prominent in the group treated with shDiPRO1/jetPEI® relative to siDiPRO1/jetPEI® (Fig. 4E). This was correlated with a stronger inhibiting effect on DiPRO1 gene expression (Fig. 4B). Of note, the weight maintenance of treated mice suggests an absence of significant toxicity of the DiPRO1-inhibiting nanomedecines (Fig. 4F). The antitumor activity of DiPRO1 depletion was also confirmed by morphological and immunohistochemical analyses of tumor biopsy sections at day 12 after treatment initiation (Fig. 4G). In controls, specimens were composed of malignant proliferative Ewing sarcoma consisting of large atypical polygonal eosinophilic cells associated with focal necrotic areas. At its periphery, the tumor was ill-defined with irregular nests of tumor cells invading the adipose tissue of the mouse hypoderma, indicating aggressive behavior of the tumor. Concerning the siDiPRO1/jetPEI® treated group, a similar aspect of the proliferation was observed, including again focal necrotic areas. However, the peripheral limit of the tumor exhibited a different pattern. The tumor cells were well limited by a thin regular layer of normal striated muscle cells tending to encapsulate the tumor. The same peripheral limitation of the proliferation was also observed in the shDiPRO1/jetPEI® group. The major change here was the presence of vast amounts of pink necrotic tissue suggestive of tumor destruction. These results were confirmed by TUNEL staining of the tumor biopsy sections used to assess the induction of apoptosis. The mean proportion of TUNEL positive cells/field (Fig. 4H) in the controls was 0.08%, whereas the si- and shDiPRO1/jetPEI® treated tumors had a mean proportion 3.1% and 2.3%, respectively, corresponding to a 39.4- and 28.2-fold increase. Immunostaining of the active form of caspase-3 protease revealed caspase-3 activation inside of tumor cells in both DiPRO1-depleted tumors, which was more pronounced at its periphery.

**Figure 4.**
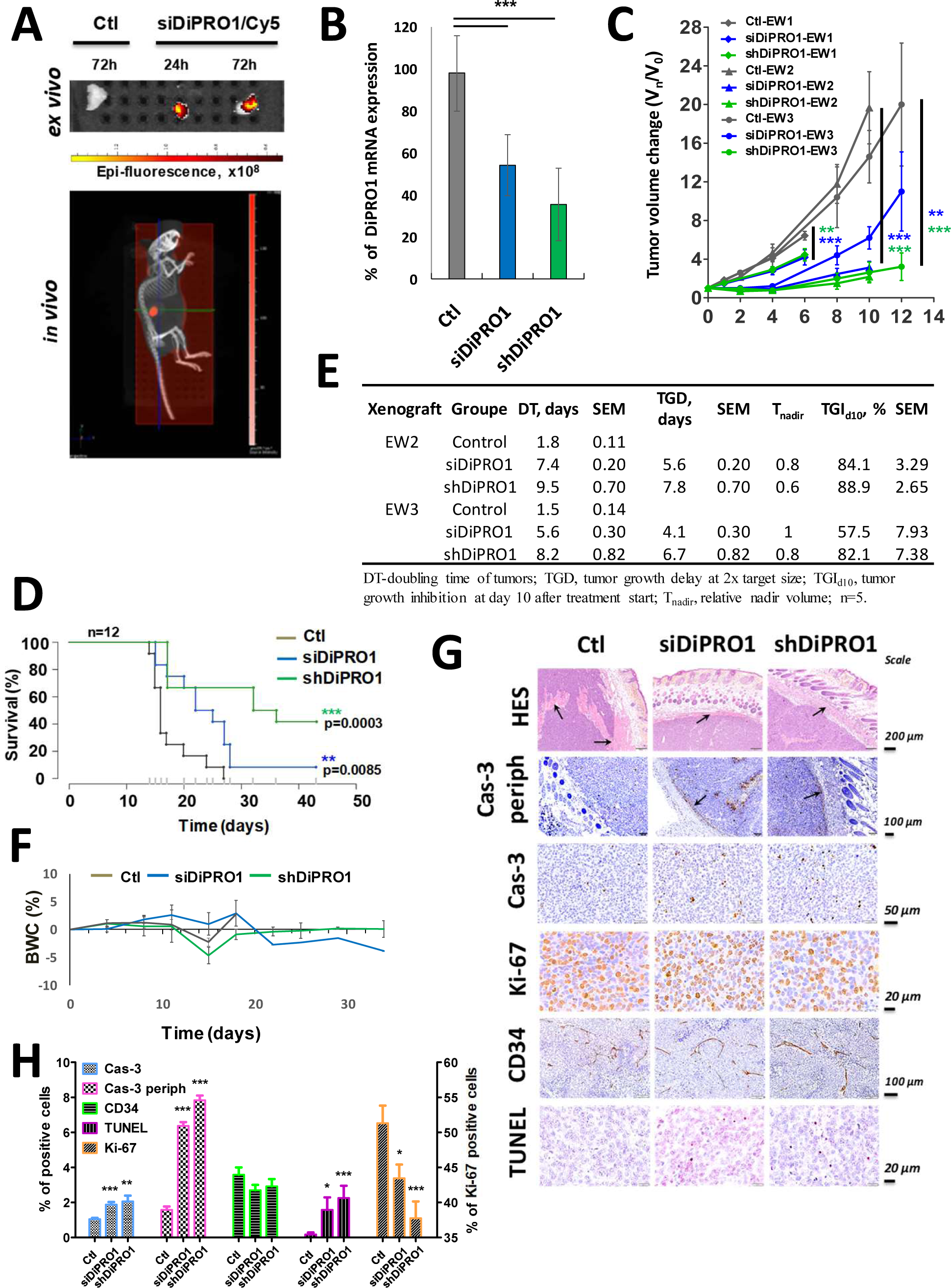
Antitumor activity of DiPRO1 inhibitors in the Ewing sarcoma (EW) subcutaneous tumor xenograft model. Nude mice received s.c. inoculations of A673 Ewing sarcoma cells and were treated with siDiPRO1/jetPEI® and shDiPRO1/jetPEI® nanocomposites or siCtl/jetPEI® scramble (Ctl) at a dose of 0.5 or 1 mg/kg/d. The initial tumor volume on the day of treatment initiation was V0=168±76 mm3 in the first (EW1; n=7 mice per group) and V0=71±24 mm3 in the second (EW2) and third (EW3) independent experiments (n=5 mice per group). **A,** Uptake by tumor cells of siDiPRO1/jetPEI®/Cy5 nanocomposites. Internalization was followed in live mice (video; lower panel) treated with a single dose (0.5 mg/kg) of Cy5-coupled anti-DiPRO1 siRNA complexes and in extracted tumors (upper panel) 24 h and 72 h after treatment. Control mice were treated with an equimolar dose of non-targeted siCtl/jetPEI® free of Cy5. **B,** DiPRO1 depletion efficiency in tumors was verified by RT-qPCR. DiPRO1 mRNA expression levels in three independent experiments were normalized to GAPDH, n=9, mean ± SD. DiPRO1 expression in control tumors was referenced to 100%. Statistical analysis was performed using the t -test. **C,** Tumor progression was compared between the indicated groups (V_n_/V_0_). Results from three independent experiments are presented. Two-way ANOVA test was used for statistical analysis at the indicated time points. **D, A**nalysis of main endpoints of antitumor activity of the si/hDiPRO1 and control nanomedicines against human Ewing sarcoma xenografts. **E,** Effect of DiPRO1 inhibition on survival of tumor-bearing mice. Kaplan-Meier curves of overall survival for 43 days (n = 12) from the day of xenograft implantation are shown. Log-Rank test (Mantel-Cox) for each pairwise comparison (treatment vs. control) was applied for statistical analysis. Results from two independent experiments (EW1 and EW2) are combined. **F,** Assessment of body weight loss (BWL) in mice bearing tumor xenografts. The results of two independent experiments (EW1 and EW2) are combined. **G,** Immunohistochemical staining of excised tumor tissues at day 12. Representative images of tumor sections stained with hematoxylin/eosin/safranin (HES) (arrows indicate striated muscle), and cells positive for internal and peripheral caspase-3 (Cas-3 periph, indicated by arrows), Ki-67, CD34 and TUNEL are shown. **H,** Quantitative analysis of positively immunostained cells (%) relative to the total number of tumor cells. Six to eleven fields were selected for counting. Necrotic fields were excluded. **C, F:** All box-plots and growth curves are displayed as means ± SEM. * P < 0.05, ** P < 0.01, *** P < 0.001, P values are shown between indicated groups and appropriate controls.

Tumors from mice receiving the si- and shDiPRO1/jetPEI®-PEI nanoparticles revealed 4.0- and 5.0-fold increases of peripheral caspase-3 expression and 1.2- and 1.4-fold decrease, respectively, of Ki-67 expression, a marker of proliferating cells. No significant difference in CD34, a marker of early hematopoietic and vascular-associated tissue, was observed at this stage of treatment. It is noteworthy that shDiPRO1/jetPEI®-PEI treatment in EW2 resulted in the survival of 60% of animals for 84 days after xenograft implantation. Histological analysis of their tumors revealed very small specimens without viable malignant cells or with a weak presence of viable tissue among a large area of necrotic tumor (Supplementary Fig. S4L).

In conclusion, nanocomposites inhibiting DiPRO1 show effective regression of tumor growth rate of Ewing sarcoma and absence of weight loss in animal models. Apoptosis-mediated tumor cell death verifies the feasibility of targeting DiPRO1 and is consistent with the results of the *in vitro* study.

### DiPRO1 upregulation in recurrent pediatric tumors. Clinical relevance

To gain further insight into the role of DiPRO1 in mesenchymal cancer development and to extend its role to other pediatric cancers, we explored whole exome and bulk RNA sequencing data from the international precision medicine MAPPYACTS trial (NCT02613962) (57). The analysis cohort included 340 patients (median age 13 years), including 37 patients with RMS and 25 patients with Ewing sarcoma, who underwent tumor biopsy/surgery for their recurrent/refractory malignancy (Supplementary Tables S1-2).

First, we performed a cross-sectional analysis of DiPRO1 expression and its downstream gene network that showed the highest fold change (FC > 1.4, pV < 0.05) in *in vitro* DiPRO1 KD experiments performed in RMS and Ewing sarcoma cells (Fig. 5A, Supplementary Fig. S5A-B). Compared with other genes, DiPRO1 showed modest expression level and expression fold change in cancers (Supplementary Fig. S5A), as the majority of KZFP (58). To elucidate the transcriptional network of DiPRO1 in patient tumor samples, we performed correlation analysis. We found a specific cluster of genes that were positively coregulated with DiPRO1 in all tumors (Fig. 5B). Interestingly, the DiPRO1-gene cluster explicitly recognizes not only the expected tumor types, namely RMS and Ewing sarcomas, but also medulloblastomas and neuroblastomas (Fig. 5C), showing the most significant upregulation of DiPRO1 in these particular tumors (Fig. 5D). Of note, the expression level of DiPRO1 was comparable between ERMS and ARMS clinical subtypes (Supplementary Fig. S5C), suggesting that its expression is not linked to the PAX3/7-FOXO1 gene fusion, a hallmark of ARMS (59). DiPRO1 and its network were then analyzed on patient-derived cancer cell lines from the Broad Institute collection. The expression pattern was consistent with that observed in primary tumors (Supplementary Fig. S5D-F, Supplementary Data S2A). Intriguingly, half of the DiPRO1 cluster genes that were upregulated in these four tumor types were identified as SOX2 targets (Fig. 5E) (60), suggesting a synergy between SOX2 and DiPRO1 to activate common downstream genes. The primary role of SOX2 is related to early neurogenesis (61), which implies a possible role of DiPRO1 in the regulation of neuromesodermal precursors and neuronal plasticity (62).

**Figure 5.**
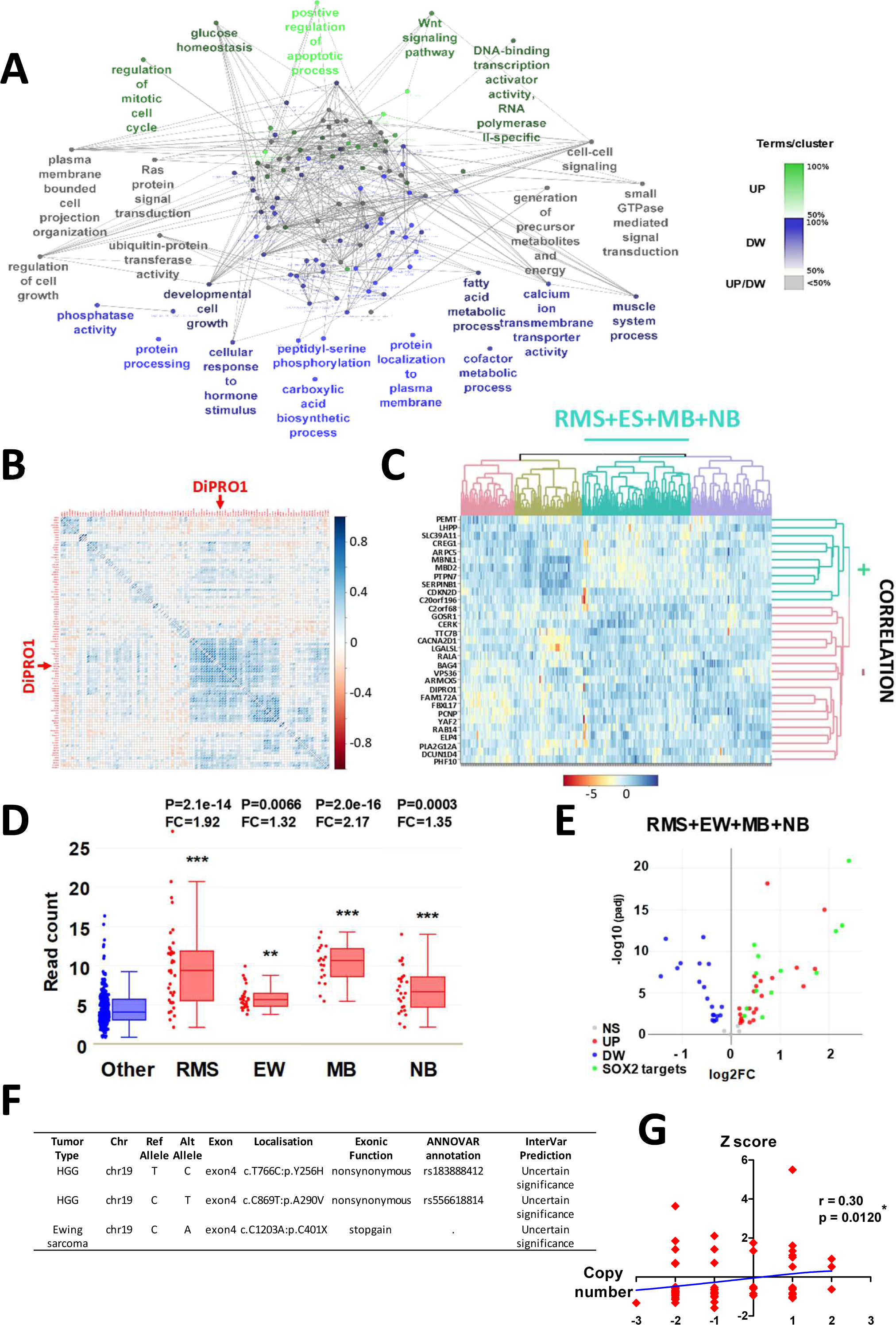
Clinical relevance of the DiPRO1 gene and its downstream targets in primary pediatric tumors. **A,** Gene ontology analysis of genes most significantly affected by DiPRO1 knockdown in both RMS and Ewing sarcoma cell models in vitro (FC=1.4, P < 0.05). The network of enriched functions of the up-regulated (n=35, green nodes) and down-regulated (n=68, blue nodes) gene groups is shown. Functions common to both groups are represented by gray nodes. Parameters were Benjamini-corrected terms, kappa score = 0.4, H. sapiens, genes in GO_BiologicalProcess. **B,** The correlation plot shows a group of genes that coexpress with DiPRO1 in pediatric tumors. **C,** The heatmap of DiPRO1 expression and its positively and negatively correlated genes distinguishes a cluster that includes RMS, Ewing sarcoma (ES), medulloblastoma (MB), and neuroblastoma (NB). Heatmap parameters: Euclidean distribution, complete clustering linkage. **D,** Boxplots of DiPRO1 overexpression in RMS (n=37), ES (n=25), MB (n=19), and NB (n=26) tumors colored in red versus other pediatric tumors (Other, n=233) colored in blue. Data are presented as median ± dispersion. **E,** SOX2 targets among DEGs commonly upregulated in RMS, ES, MB, and NB tumors are shown in the volcano plot. The cutoff threshold padj = 0.05. Regulator analysis was performed using the Cytoscape iRegulon application. The SOX2 motif (tfdimers-MD00490) showed the highest normalized enrichment score (NES=6.349) within 14 DEG-promoters. **F,** Single nucleotide variations (SNVs) of DiPRO1/ZNF555 in pediatric tumors. The analysis was performed by exploring whole exome sequencing data on pediatric tumor specimens. **G,** Integrative analysis of the association between DiPRO1 copy nimbler variation (CNV) and differential gene expression. XY scatter plot shows the relationship between Z score (y -axis) and copy number value (x-axis) in 101 pediatric tumor samples. The correlation curve is shown in blue. Spearman’s correlation coefficient (r) of the nonlinear regression fit and p-value with Gaussian approximation between copy number and mRNA expression are indicated. **B-E:** RNA-seq data of primary tumor biopsies were processed using DEBrowser, DESeq2 (TMM normalization, parametric fit, LRT test) and ggplot2 R package. RMS: rhabdomyosarcoma; ES: Ewing sarcoma. MB: medulloblastoma, NB: neuroblastoma, HGG: high-grade glioma. ns: non-significant P > 0.05, *P > 0.05; **P > 0.01; *** P < 0.001. P-values are shown between indicated tumor type and other samples.

We next addressed whether abnormal DiPRO1 expression could be caused by somatic mutations or copy number variations. According to our study, no somatic DiPRO1 mutations were detected in RMS patients, whereas three single nucleotide variants (SNVs) were found in exon 4 of two patients, an exonic stop-gain mutation (exon4: c.1203 C→A) in a Ewing sarcoma and two synonymous SNVs (exon4: c.766 T→C and c.869 C→T) in a high grade glioma (Fig. 5F, Supplementary Data 2B). These variations of uncertain significance (InterVar prediction) did not alter DiPRO1 gene expression compared with other samples within the same cancer type and can therefore be considered as non-functional. We additionally explored external COSMIC (Catalogue of Somatic Mutations In Cancer) data (63) for fusions, translocations, and mutations of the DiPRO1 gene and found no alterations in its coding sequence in human cancer, which was consistent with our results. Integrated analysis of copy number variations (CNVs) and differential gene expression in pediatric cancers showed that DiPRO1 DNA copy number was altered in 29.4% (101 of 343) of tumor samples, showing a weak correlation (r = 0.30) (64) with DiPRO1 gene expression level (Fig. 5G, Supplementary Data 2C). The frequency of high copy numbers of the DiPRO1 gene with high gene expression was remarkably low (2.3%) compared with conventional oncogenes (64, 65), albeit among these eight samples, three were from RMS and one from medulloblastoma.

Overall, pan-cancer analysis of recurrent tumor biopsies argues for DiPRO1 targeting in mesenchymal tumors, showing increased expression levels of DiPRO1 together with SOX2-linked genes. Based on mutational analysis, DiPRO1 does not exhibit the properties of classical oncogenes.

### DiPRO1 physically interacts with TIF1B and UHRF1

In previous sections, we showed that DiPRO1 can bind promoter, gene body and intergenic regions enriched with retroviral repeat elements. DiPRO1 regulates large-scale transcriptional network and its downregulations leads to cancer cell death. To elucidate the molecular mechanism by which DiPRO1 regulates its transcriptional network, we sought to determine the partners with which DiPRO1 interacts. To this aim, the pDiPRO1 vector expressing the DiPRO1-YFP protein with FLAG and HA tags was transfected into RMS TE671 cells and the expression of exogenous DiPRO1 mRNA and protein was verified (Fig. 6 A-B and Methods section).

**Figure 6.**
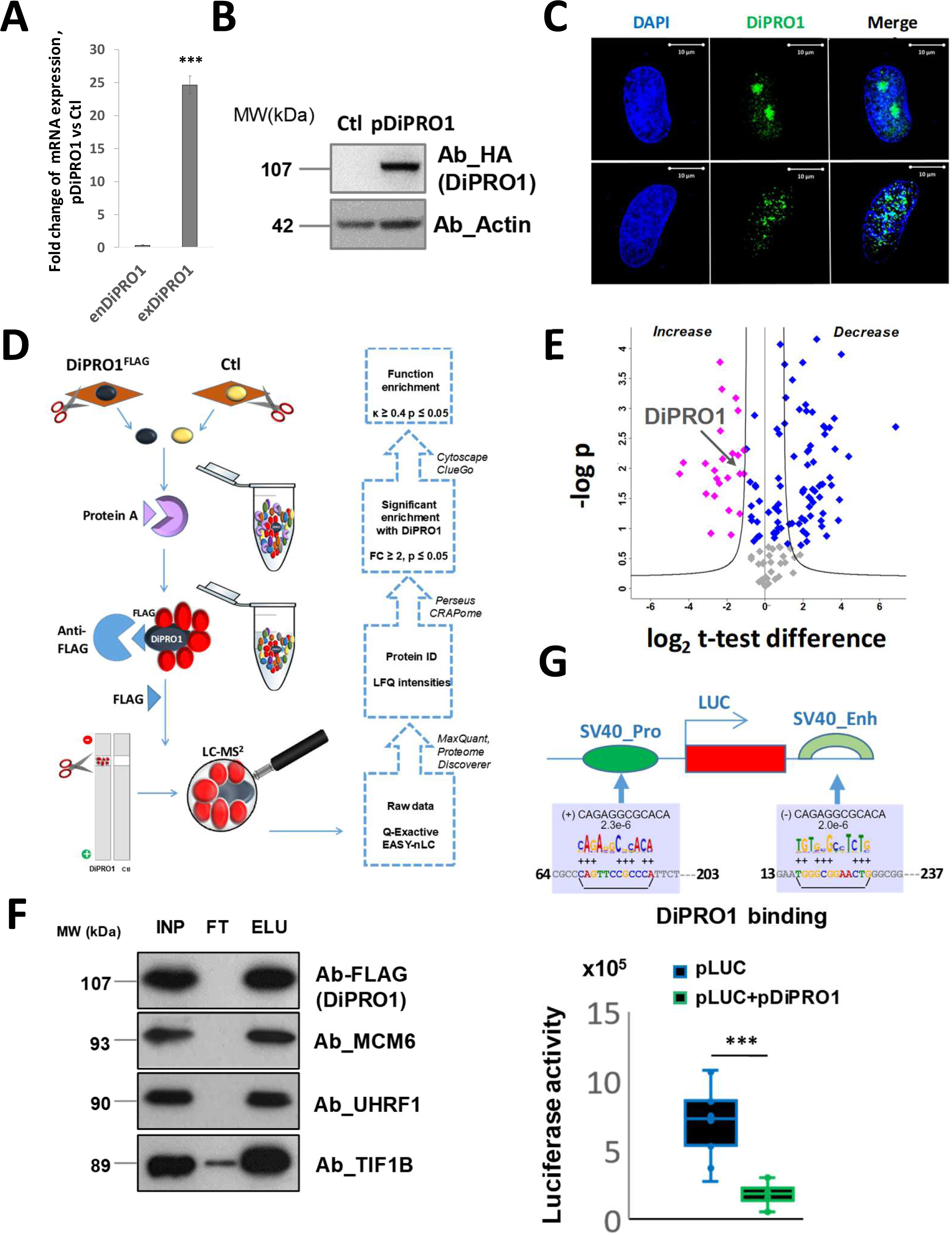
DiPRO1 transcriptional activity and interaction partners. **A-G,** The RMS (TE671) were transiently transfected with a retrovirus vector expressing an ORF of DiPRO1 and FLAG, 6-His and HA tags (pDiPRO1^FLAG, 6-His, HA^). The efficiency of exogenous DiPRO1 overexpression was verified by mRNA expression analysis **(A)** and western blotting **(B)** 24h post-transfection. The RT-qPCR analysis was performed using the primers for exogenous (exDiPRO1) and endogenous DiPRO1 (enDiPRO1) cDNA. The results of qRT-PCR were normalized to the mRNA expression of GAPDH gene and represent three independent total RNA extractions measured in triplicate, mean ±SD. Ct=45 was set up for exDiPRO1 ORF expression in control cells. t-test was used for statistical analyses. ***P < 0.001 indicates the difference between endogenous and exogenous DiPRO1 expression. Immunoblotting assays were performed with whole cell extracts from RMS cells overexpressing DiPRO1 (pDiPRO1) or vector control (Ctl). The DiPRO1 protein level was analyzed using anti-HA antibody. The Actin was served as an internal control. **C,** Confocal microscopy analysis revealed the expression of DiPRO1-fused YFP. Scale bar: 10 µm. **D,** Sample overview and methodological workflow. Nuclei extracted from RMS cells stably expressing pDiPRO1^FLAG^ and their isogenic counterparts, lacking expression of the bait protein (Ctl), were pre-cleared with protein A beads, immunopurified with an anti-FLAG affinity column, and eluted by adding FLAG-competitor. Eluates were resolved by SDS-PAGE and the corresponding protein bands were excised, processed, and analyzed by mass spectrometry (LC-MS2). IDs—identifications; LFQ—label-free quantification. **E,** The volcano plot represents proteins that were differentially enriched (blue dots) between groups of RMS cells expressing DiPRO1 (pDiPRO1) and parental cells (Ctl). Pink dots correspond to proteins enriched in the DiPRO1 complex. Two-sample t-test was applied with threshold s0 = 2 and FDR = 0.05. Data represent independent biological replicates (n=2-3). **F,** Identification of MCM6, UHRF1 and TIF1B^FLAG^ proteins in DiPRO1-precipitates by Western blot. The nuclear extracts from pDiPRO1 RMS cells were immunoprecipitated with antibodies against FLAG followed by immunoblotting with antibodies against FLAG, MCM6, UHRF1 and TIF1B. INP: Input, FT: Flow-Through and ELU: DiPRO1-immunoprecipitated elution. **G,** Overexpression of DiPRO1 inhibits the activity of the gene enhancer and promoter. The pDiPRO1 was cotransfected with a set of luciferase reporter constructs (pLUC) containing the SV40 promoter (SV40pro) alone or with the SV40 enhancer (SV40e). Data were scaled and the luciferase signal of the pGL3 base vector was extracted. Results represent four experiments performed in triplicate, mean ± SD, n=10-12. The t-test for paired differences between transfections with and without pDiPRO1 was performed, ***P < 0.001.

Consistent with its predicted role in transcriptional regulation, DiPRO1 is strictly localized in the nucleus (Fig. 6C). Accordingly, the affinity purification coupled with mass spectroscopy (MS) was performed using nuclear fraction (Fig. 6D). Pulldown analysis revealed twenty-four proteins (cutoff s0 = 2 and FDR = 0.05) enriched by the FLAG-tagged DiPRO1 bait protein in the nuclei of RMS cells expressing DiPRO1 (Fig. 6E). The putative DiPRO1 interacting candidates were then filtered on the basis of 1) at least two identified peptides, 2) significantly higher intensity of the identified peptides in DiPRO1-expressing cells compared with controls, 3) low CRAPome score, and 4) nuclear localization. Among ten selected proteins (Supplementary Data S3), the KRAB co-repressor TIF1B/KAP1 and UHRF1/NP95 appeared in the precipitated complex with DiPRO1. It should be noted that, in addition to gene expression regulation, TIF1B and UHRF1 play a critical role in the epigenetic silencing of retroviral elements (66–68). In addition, four proteins of the hexameric mini-chromosome maintenance complex (MCM) were found enriched in the DiPRO1 protein complex, confirming the involvement of DiPRO1 in cell cycle control that we previously demonstrated. The physical interaction of DiPRO1 with TIF1B, UHRF1 and MCM6 was validated by anti-FLAG immunoprecipitation (IP) and Western blot analysis in RMS cells (Fig. 6F).

We previously reported that the inhibition of DiPRO1 led to the partial loss of β-satellite enhancer activity (4qAe), indicating a transcriptional activator characteristic of DiPRO1 (17). Based on the properties of the interacting module proteins, TIF1B (69) and UHRF1 (70), we hypothesized that DiPRO1 might be a transcriptional repressor. According to the motif prediction, DiPRO1 may have binding sites in the SV40 promoter and enhancer and thus may regulate gene transcription through viral *cis*-elements. To verify this, we applied luciferase reporter assays using the pGL3 reporter vector containing the SV40 promoter (Pro) and enhancer (Enh). The base vector without promoter and enhancer was used for normalization. The vectors were transfected to RMS cells alone or co-transfected with pDiPRO. The results of the chemiluminescence signal quantification showed that DiPRO1 strongly reduced reporter gene expression (Fig. 6G), indicating that DiPRO1 could repress enhancer/promoter-mediated gene expression.

### DiPRO1 globally regulates gene repression and RE silencing through CGI methylation

Our results in RMS cells established the physical interaction of DiPRO1 with TIF1B and UHRF1. Both proteins can participate in the methylation regulation of CpG islands (CGI) (67, 71). While TIF1B plays the role of a scaffold to recruit DNA methyltransferas (DNMTs) (39), UHRF can bind directly to methylated DNA via its SRA domain (72). Thus, we hypothesized that DiPRO1 may contribute to gene and RE regulation via CGI methylation. To characterize this relationship, we applied the methylated CpG island recovery assay sequencing (MIRA-seq) to RMS (TE671) cells transduced with shDiPRO1- and shCtl-expressing vectors 48 hours following infection. Consistent with previous studies, the upstream CpG regions of the PAX3 (73) and MYOD1 (74) genes were methylated, while those of the fibroblast growth factor receptor 1 (FGFR1) (75) and JUP (γ-catenin) (76) genes were unmethylated in control RMS cells (Supplementary Fig. S6A). Further analysis revealed a methylated genome size of 5.5 Mb and 14.3 Mb in shDiPRO1 and shCtl, respectively, which was enriched compared to the corresponding inputs (INP) (Fig. 7A). This yielded 3,984 differentially methylated unique CGIs (DMCs) in shDiPRO cells, which were designated as hypermethylated DMCs, and 11,283 DMCs in control cells, which were designated as hypomethylated DMCs (Supplementary Data S4). These observations indicate that at the genome-wide level, DiPRO1 is involved in CGI methylation, and its downregulation leads to CGI hypomethylation. Genomic distribution analysis showed that in both shDiPRO1 and shCtl, about 60% of differential methylation occurred in the core gene regions, represented by a population of long housekeeping-CGIs (hk-DMCs) of an average length of 777 bp, and a population of short repeat-associated CGIs (rep-DMCs) of an average length of 309 bp (Fig. 7B, Supplementary Fig. S6B-C). The remaining portion (40%) came from intergenic regions, mainly comprising rep-DMCs. Globally, rep-DMCs were mostly hypomethylated and constituted the majority (65.5%) of all hypomethylated DMCs (Fig. 7C). Our previous analysis showed that DiPRO1 can bind directly different classes of RE. We then addressed which classes of repeats are affected by DiPRO1 KD. Although hypermethylation mainly took place within SINE repeats (92%) (Supplementary Fig. S6D), hypomethylation was detected in different classes of repeats (SINE, LINE, LTR/ERV) (Fig. 7D), which was consistent with the DiPRO1 binding analysis. Accordingly, we can speculate that the binding of DiPRO1 to REs may contribute to their silencing by methylation.

**Figure 7.**
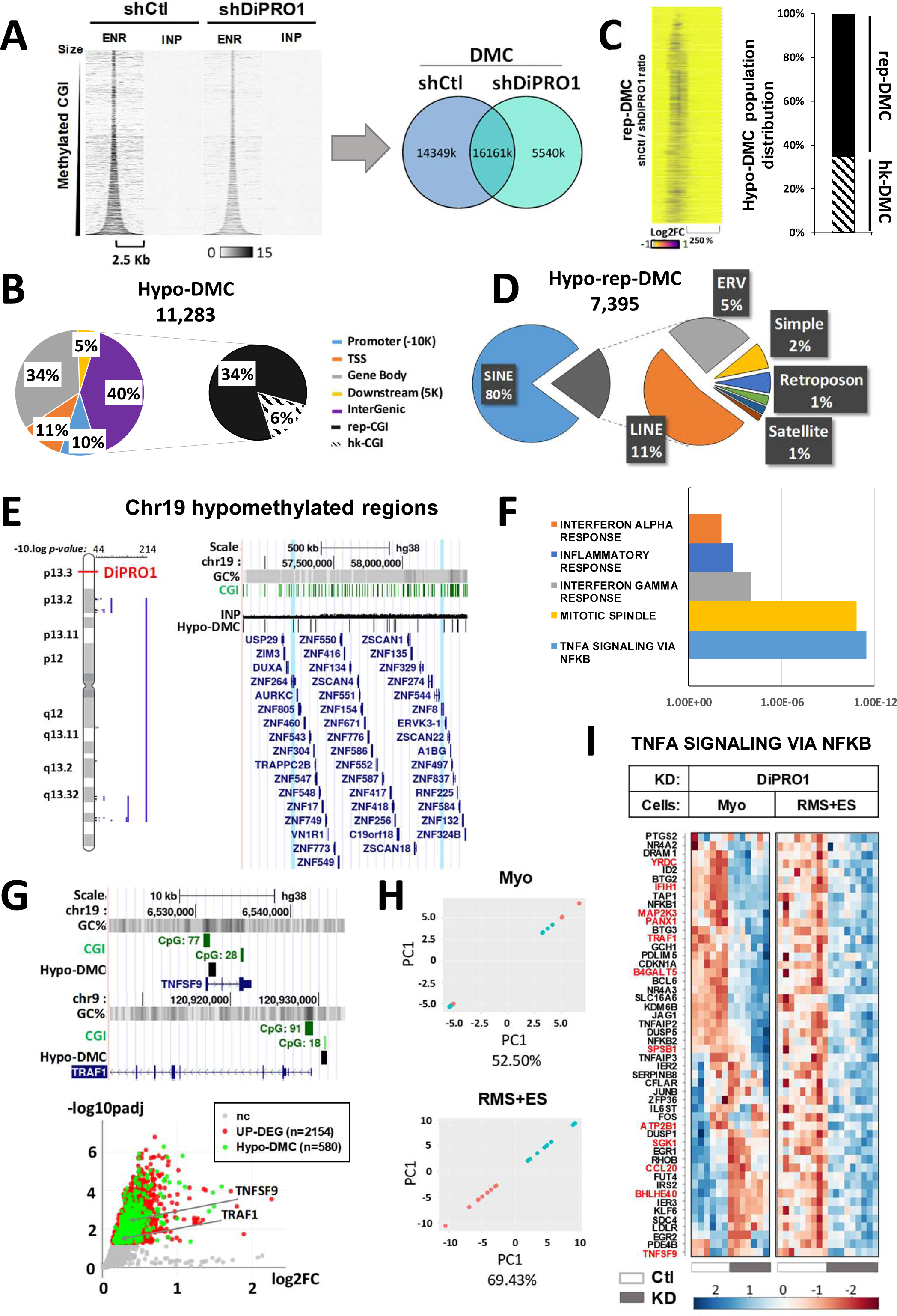
Methylation changes in different CpG island populations related to DiPRO1 inhibition in RMS cells. **A-G**, Methylation using MIRA-seq was analyzed in DNA samples from TE671 cells. Cells expressing a DiPRO1-targeting shRNA (shDiPRO1) were compared with control cells expressing a non-targeting shRNA (shCtl). Signals from methylation-enriched DNA (ENR) were normalized to the unenriched input DNA (INP). **A,** Heatmaps (top) displaying methylation levels in size-sorted CGIs in shDiPRO1 and shCtl and corresponding inputs. The bar reflects the signal intensity. Y-axis: DNA fragments per 1M reads per 1K. X axis: 2.5 Kb surrounding each region. Venn diagram (bottom) showing the overlap in bp for the size of shDiPRO1 and shCtl regions in the CGIs. **B**, Genomic distribution of hypomethylated DMC is shown as % of total number of DMC. **C,** The rep-CGI population abundance within hypomethylated DMC. Ratiometric heatmap (left) of rep-DMC showing log2 fold change in two signals (shCtl/sh DiPRO1). The color bar reflects the signal intensity. Y-axis: DNA fragments per 1M reads per 1K. Distribution (%) of hk-DMC and rep-DMC populations (fight). **D,** DNA repeat class distribution within DiPRO1-linked hypomethylated DMC regions. **E,** Hypomethylated DMC signals are overrepresented in chromosome 19. The analysis was performed using the Positional Gene Enrichment (PGE) tool. **F,** Hallmark enrichment of genes associated with hypomethylated DMC identified in DiPRO1 KD cells. **G**, The TNFSF9 and TRAF1 genes were hypomethylated in TSS regions and upregulated in DiPRO1 KD cells. CGI, GC (%) and hypomethylated DMC tracks over the TNFSF9 and TRAF1 loci (top). Volcano plot (bottom) shows upregulated DEGs (red dots, p<0.05), which are hypomethylated in DiPRO1 KD cells (green dots). **H**, PCA analysis of DEGs of the TNFA signalling via NFKB pathway of DiPRO1 KD in RMS and Ewing sarcoma cells and myoblasts. **I**, Heatmap expression of the TNFA signalling via NFKB pathway gene set across control and DiPRO1 KD samples of RMS and Ewing sarcoma cells and myoblasts. Clustering analysis of Euclidean distribution, complete linkage **G-I**: Global transcriptome analysis by microarray was implemented using total mRNA extracted from RMS and Ewing sarcoma cell lines: with DiPRO1 knockdown (shDiPRO1) and corresponding controls (shCtl). Three to six separate extractions from two independent experiments were performed to collect the replicates (n = 9 per condition). Differentially expressed genes in shDiPRO1-cells versus corresponding controls (P < 0.05) were analyzed using the Limma R package. CGI, CpG islands according to Gardiner-Garden and Frommer criteria; DMC, differentially methylated CGI; hk-DMC, housekeeping DMC w/o repetitive elements; rep-DMC, DMC with repetitive elements.

To establish the functional significance of methylation changes resulting from DiPRO1-KD, we analyzed the correlation between methylation and gene expression. We focused on hypomethylation precisely as it was the predominant event in DiPRO1 KD. CpG methylation is traditionally thought to be associated with gene repression (77). Consequently, hypomethylated DMCs were confronted with upregulated DEGs in DiPRO1 KD and control cells. About 14.6% of DMCs showed a correlation with the gene transcription changes in RMS cells (Supplementary Fig. S6E). Interestingly, they coincided with both hk-DMCs and rep-DMCs, suggesting that the population of rep-CGIs may contribute to gene expression. This is in accordance with previous work on epigenetic regulation of DNA repeat silencing by KZNP (10, 78) and the contribution of DNA repeats to the regulation of gene expression (79). Upon functional analysis based on KEGG pathways we observed the enrichment of pathways in cancer, virus infection, axon guidance, neurodegeneration and endocytosis (Supplementary Fig. S6F). Upregulation of positive regulators of apoptosis, the CASP2, CASP5, CDKN1A, MAP3K5/ASK1, and PYCARD genes (55), was associated with hypomethylation of proximal CGIs. Interestingly, the huge army of ZFPs and KZFP regrouped in the viral infection pathway was overrepresented in chromosome 19, according to the Positional Gene Enrichment (PGE) analysis (80) (Fig. 7E). It appears that chromosome 19 was particularly affected by DiPRO1-linked methylation. According to the GSEA/MSigDB database of hallmarks (51), we observed that DiPRO1 KD induced the hypomethylation of genes involved in inflammation, including tumor necrosis factor-α (TNF-α) signaling via NF-kB (Fig. 7F). Among them the adaptor protein TRAF1 and the TNF-receptor TNFRSF9, key components of TNF-α signaling, were hypomethylated and upregulated in DiPRO1 KD (Fig. 7G). Their interaction is important for the transmission of signals leading to the activation of NF-kappaB, an anti-apoptotic factor. At the same time, TRAF1 is a specific target of activated caspases for apoptosis induction in cancer cells, playing the role of a link between caspases and TNF-receptors (81). It has been reported that increased TNFRSF9 levels lead to suppression of tumor progression in breast cancer (82), suggesting its possible contribution to the antitumor activity of DiPRO1. Principal Component analysis (PCA) shows that the gene signature of TNF-α signaling via NF-kB segregates the control from the DiPRO1 KD in RMS and ES cells, but not in myoblasts (Fig. 7I). The transcriptome analysis demonstrates a clear shift in the expression pattern of these genes from low expression in control cells to higher expression in DiPRO1 KD cells (Fig. 7H).

Thus, our results support the hypothesis that DiPRO1 is a regulator of CGI methylation. Its loss leads to preferential demethylation of CGIs associated with both gene core regions and DNA repeats, which contributes to the deregulation of apoptotic genes as well as genes involved in viral infection and inflammation.

### DiPRO1 and SIX1 crosstalk in mesenchymal cancer

Because DiPRO1 shares common targets with SIX1 and regulates the myogenic transcriptional program in normal muscle cells, we wondered whether the DiPRO1 and SIX1 gene regulatory network is common to mesenchymal cancer cells. Using SIX1 targets from harmonized ChIP-seq, ChIP-exo, DNase-seq, MNase-seq, ATAC-seq and RNA-seq data (83), we showed that 12.4% of promoter regions bound by SIX1 overlap with DMCs in DiPRO1 KD with predominance in hypomethylated regions (10.0%) (Fig. 8A). This supports the possibility of DiPRO1-linked methylation in SIX1 binding regions. Using the recently published dataset of SIX1 KD in human RMS (32), we compared its transcriptional network with that of DiPRO1 KD. The overlapping pattern of upregulated genes in DiPRO1 KD corresponds to 8.2% in RMS cells and 10.8% in myoblasts. As expected, these genes were enriched in the main hallmarks of myogenesis (Fig. 8B). As illustrated in the Venn diagram (Fig. 8C), the myogenic signature of SIX1 KD cells closely resembled that of DiPRO1 KD myoblasts, including upregulation of MYOG, ACTA1, and the troponin subunits TNNC1/2, TNNT3, and TNNI1. In RMS cells with SIX1 KD and DiPRO1 KD, we found commonly increased levels of the embryonic MYH3 gene, early myogenic genes MEF2a/c, and contractile muscle genes MYOZ1, MYOM1, MYL1, and TNNT2, showing that both proteins in RMS slow down the differentiation program. Furthermore, depletion of DiPRO1 induced upregulation of additional myogenic genes DMD, TCAP, and MYF6/MRF4, which are important players in muscle differentiation and contraction. A very limited number of myogenic genes were upregulated in a pooled pattern across ES and RMS cells upon DiPRO1 KD. While MYOM2 is a striated muscle specific gene, encoding a sarcomeric protein, the MAPRE3/EB3, AK1, ABLIM1, BHLHE40/DEC1 genes share neuronal and muscle functions. KEGG-based functional analysis revealed enrichment of nervous system pathways in DEGs that were hypomethylated and upregulated upon DiPRO1 KD (Supplementary Fig. S6F).

**Figure 8.**
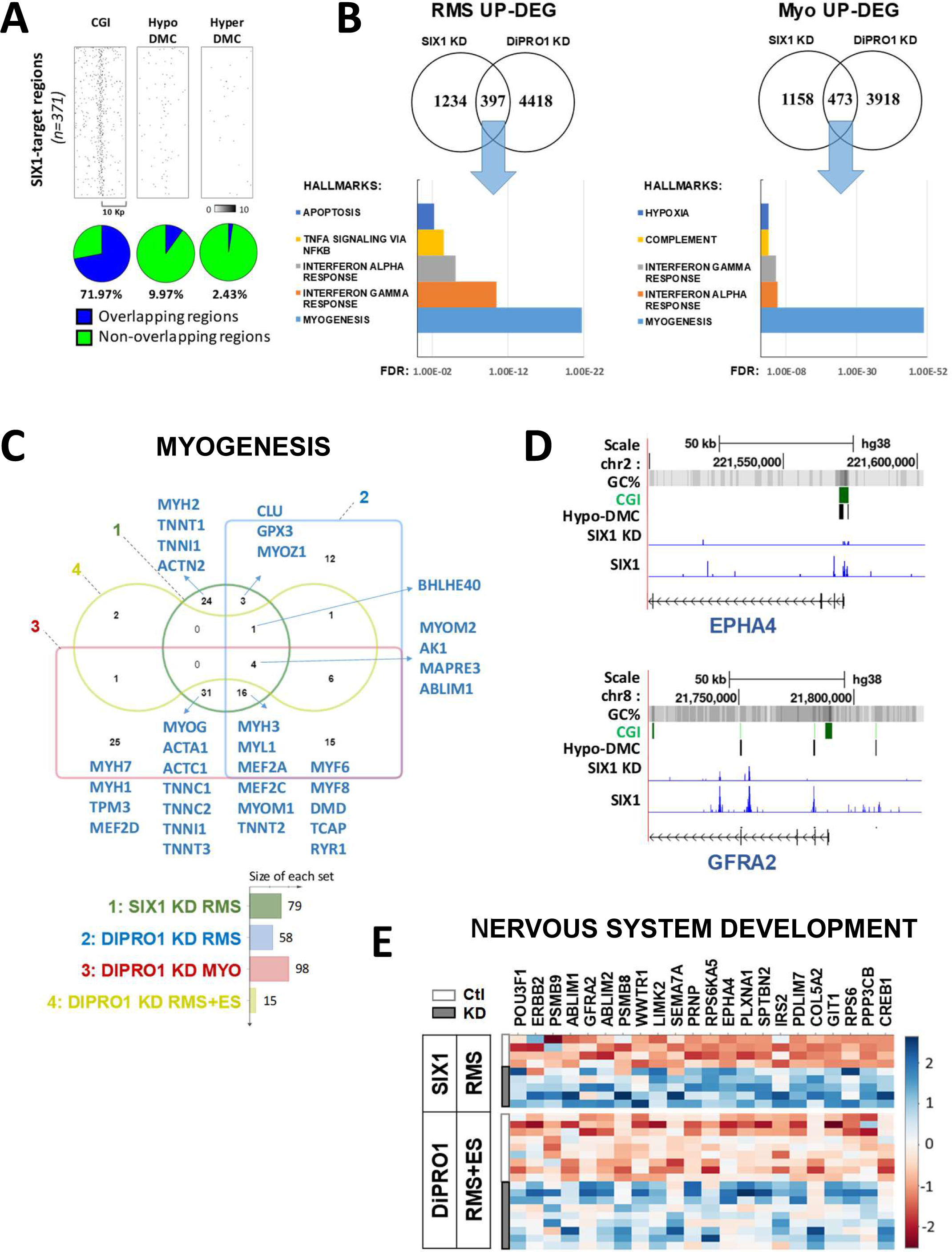
DiPRO1 and SIX1 KD common signature. **A-G**, Heatmaps (top) displaying SIX1 target signals from the GSEA data collection within CGIs of hg38, hypo- and hypermethylated DMC in DiPRO1 KD RMS. The bar reflects the signal intensity. Y-axis: DNA fragments per 1M reads per 1K. X axis: 10 Kb surrounding each region. Venn diagram (bottom) showing the overlapping regions. **B**, Hallmark enrichment analysis of shared upregulated DEGs in SIX1 KD and DiPRO1 KD in RMS (left) and myoblasts (right) **C,** Venn diagramm of the myogenesis gene set shows a joint signature of SIX1 KD and DiPRO1 KD. **D,** The tracks of DiPRO1-linked hypomethylated DMCs, SIX1 control and KD ChIP-seq regions over the EPHA4 and GFRA2 loci. **E,** Heatmap of common expression gene set of the nervous system development KEGG pathway across DiPRO1 KD and SIX1 KD samples and appropriate controls of RMS and Ewing sarcoma (ES) cells. Clustering analysis of Euclidean distribution, complete linkage, p<0.05. **A,D:** Methylation using MIRA-seq was analyzed in DNA samples from TE671 cells. Cells expressing a DiPRO1-targeting shRNA (shDiPRO1) were compared with control cells expressing a non-targeting shRNA (shCtl). Signals from methylation-enriched DNA (ENR) were normalized to the unenriched input DNA (INP). DMC, differentially methylated CGI. **B, C, E**: Global transcriptome analysis by microarray was implemented using total mRNA extracted from RMS and Ewing sarcoma cell lines: with DiPRO1 knockdown (shDiPRO1) and corresponding controls (shCtl). Three to six separate extractions from two independent experiments were performed to collect the replicates (n = 9 per condition). Differentially expressed genes in shDiPRO1-cells versus corresponding controls (P < 0.05) were analyzed using the Limma package. **B-E:** The RNA-seq and ChIP-seq dataset of SIX1 KD in RMS cells were retrieved from GSE173155. The RNA-seq data were processed using DEBrowser, DESeq2 (TMM normalization, local fit, LRT test). The ChIP-seq data were uploaded by IGV browser. RMS: rhabdomyosarcoma cells, ES: Ewing sarcoma cells.

Besides its role in myogenesis, SIX1 also functions as a neuronal fate activator in cooperation with SOX2 in mice (84, 85) and a neuronal fate inhibitor in zebrafish (86). RMS cells include the subpopulations of neural progenitors, along with cells expressing muscle developmental features (87). We therefore hypothesized that DiPRO1 and SIX1 might cooperate in sharing the neurogenic network. Accordingly, we further inspected the genes involved in the pathways of nervous system development (KEGG). As expected, the analysis revealed 22 common genes in RMSs and Ewing sarcomas that were upregulated by SIX1 or DiPRO1 KD. Among them, the genes EPHA4 (88) and GFRA2 (89), were hypomethylated in the promoter regions upon DiPRO1 KD and this coincided with the loss of SIX1 binding in SIX1 KD cells. Therefore, these results demonstrate that DiPRO1 may repress neuromuscular development by operating through shared SIX1 targets.

## DISCUSSION

We provide evidence that DiPRO1 is a transcriptional and epigenetic regulator that compromises normal and malignant cell fate in a selective manner. Functional modulation of gene expression demonstrates that DiPRO1 is a positive regulator of proliferation and a negative regulator of differentiation of human myoblasts whereas it is a positive regulator of proliferation and prevents apoptosis in cancer cells. Besides discrete expression alterations, a feature of KRAB zinc finger proteins (58), DiPRO1 inhibition is toxic to RMS and Ewing sarcoma cancer cells, suggesting that cancer cells require DiPRO1 for their survival. DiPRO1 is not a classical oncogene, its mutations are rarely observed in cancer. Targeting rarely mutated genes that are conserved by cancer cells during clonal evolution has been reported as an alternative strategy for cancer therapy (90), supporting our hypothesis of defeating cancer cells by inhibiting DiPRO1.

Previously, we reported that DiPRO1 binds the 4q35e enhancer of beta-satellite repeat region and its inhibition results in gene repression, suggesting the activator property of DiPRO1 (17). Here, we demonstrate that DiPRO1 presents binding sites in the SV40 enhancer and promoter and can inhibit gene expression. DiPRO1 binds also to DNA regions where it may contribute to CpG methylation, associated with heterochromatin. At the same time, it could be associated with the transcription at promoter and enhancer regions in active chromatin. We note that 42.6% of the genes associated with DiPRO1 binding sites in their vicinity are epigenetically activated and 22.5% are repressed by DiPRO1 KD, suggesting that DiPRO1 may exhibit a dual transcriptional role (supplemental Figure S7). This could probably related to a combinatorial recruitment of corepressors or coactivators, involved in the gene transcriptional control (15). Similar to the vast majority of KZFP (10, 13), DiPRO1 interacts with the co-repressor TIF1B, also known as KAP1. Additionally, UHRF1 ubiquitin ligase also known as NP95 was found in the DiPRO1-complex, similar to KZFP ZFP57 system (82). TIF1B and UHRF1 are classically considered to promote histone- and DNA-dependent epigenetic gene silencing (91, 92) and this may contribute to the repressive property of DiPRO1. Therefore, it remains to be determined how DiPRO1 may activate gene expression, and the positive cofactors recruited to mediate it. It has been reported a bivalent nature of KAP1 in gene regulation (93). The mechanisms by which KAP1 can activate transcription are nevertheless not fully understood. A model involves a differential phosphorylation of KAP1 that leads to release of corepressors and activation of MyoD/Mef2D promoters in myogenic cells (93). Whether this mechanism participates to the functions of DiPRO1 observed in RMS remains to be established.

One theory of carcinogenesis (94) explaining cancer development through epigenetic cell reprogramming has important implications for finding new strategies for cancer therapy (95). The DiPRO1 interaction partners UHRF1 and TIF1B have been proposed as targets for cancer therapy (96–98). Both of them can contribute to the DNA methylation recruiting mainly DNMT1 (71, 99). UHRF1 is essential to maintain CGI methylation and associated gene silencing in cancer (97). While UHRF1 can recognize hemi-methylated CGI via its SRA domain, TIF1B does not have a DNA-binding domain and thus requires another protein partner to be recruited to the genome. According to this cooperation, one of the mechanisms, by which DiPRO1 can attenuate gene expression, is associated with methylation. The disruption of DiPRO1 cooperation with CGI methylation regulators lead to massive CGI demethylation including epigenetic awakening of RE in cancer cells. Considered for decades as “junk”, DNA repeat *cis*-elements could influence the oncogene and tumor suppressor gene transcription (100). Some of them contribute to the heterochromatin integrity protecting the DNA from damage (14). Hence, it is tempting to assume that DNA repeats could constitute an “epigenetic barrier” for an anticancer response. Human endogenous retrovirus (HERV) activation in tumors drives the synchronized elevation of KZFP expression, presumably leading to tumor suppression (101). In agreement with this, DiPRO1 loss induced a massive activation of ZFP, including a large amount of KZFP, mimicking a virus infection response. ZFP of vertebrates evolutionally developed a host defense to virus infection mediated by viral ZFP (102). In addition to ZFP, the early events of DiPRO1 KD are associated with CGI demethylation-linked gene activation, which are involved in inflammation processes, mainly via TNF-α signaling. In relation with virus infection, TNF-α signaling is an important driver of host antiviral defense leading to cell death and tissue destruction (103–105). Among them the adaptor protein TRAF1 and the TNF-R receptor TNFRSF9 were demethylated and overexpressed. Their interaction is important to transduce the non-apoptotic signals leading to activation of NF-kappaB. At the same time, TRAF1 is a specific target of activated caspases for the apoptosis induction in cancer cells playing a role as a link between caspases and the TNF receptors (76). Together with upregulation of pro-inflammatory NFKB1 and NFKB2 genes and inflammatory caspases CASP4 and CASP5, DiPRO1 KD induces the expression of the initiator caspase CASP8, leading to the activation of CASP3, a final “executioner” of apoptosis (48). Nonetheless, we do not exclude other relevant forms of cell death, such as necroptosis, necrosis, or pyroptosis (48). To summarize, early events of disruption of the DiPRO1-repressive complex in cancer cells lead to mimicking the defense against viral infection and tuning the cell decision from inflammation to death.

In our study we demonstrate that DiPRO1 can share known direct targets of SIX1 and maintain their methylation, probably interfering with further SIX1 binding. SIX1 is an upstream regulator of myogenesis (18) and epithelial mesenchymal transition (19). It can also participate in the neuronal developmental program (85, 86). Several studies have reported a role of SIX1 in oncogenesis (32,106,107). It can promote the tumor growth, invasion and metastasis. In RMS, SIX1 preserves the undifferentiating state of tumor cells and SIX1 KD leads to myogenic differentiation and impaired tumor growth (32). In our study, DiPRO1 maintains suppressed muscle differentiation program in myoblasts and RMS cells. Loss of SIX1 or DiPRO1 leads to common myogenic consequences. During mouse myogenesis, murine Six1 shows a feedback regulatory loop with Pax3 and is likely to be upstream positive regulator of Pax3 (18,108,109). The myogenic alterations due to the mutation of Six1 resemble with those of Pax3 mutants (110, 111). In myoblasts and RMS, DiPRO1 positively controls the PAX3 gene expression, and PAX3 is required for RMS cell viability (112).

RMS is the most common soft tissue sarcoma in children and adolescent, considered as an aggressive and highly malignant cancer (113). Ewing sarcoma is a small round cell tumor, highly malignant and poorly differentiated that is currently the second most common malignant bone tumor in children (114). It can also develop in the extraskeletal soft tissues, including muscles (115). The shared molecular and functional changes of early events of DiPRO1 inhibition in these cancer cells led to cell cycle deregulation, differentiation and caspase-mediated cell death. Despite heterogeneity, RMS and Ewing sarcoma are considered as being derived from primitive mesenchymal stem cells (113, 114) and have been currently classified as mesenchymal tumors in the 2016 World Health Organization classification (46), supporting a possibility of common mechanism of DiPRO1 action in these cancer cells. Cell fate mapping of RMS tumors shows presence of subpopulations of neural and muscle mesenchymal progenitors that can make both muscle and osteogenic cells, along with cells expressing muscle developmental features (87). RMS can be trans-reprogramed to the neuronal phenotype resembling Ewing tumors (49) by introducing EWS-FLI1 fusion gene. Therefore, both tumors could be commonly targetable. Along with myogenic differentiation regulators, DiPRO1 can control neuronal development in RMS and Ewing sarcoma through SIX1 targets. Among them ephrin receptor EPHA4 (88), actin binding LIM-proteins ABLIM1 and ABLIM2 (116), semaphorin SEMA7A (117) and glial neurotrophic factor GFRA2 (118) genes, which are involved in axon guidance, are commonly repressed in cancer cells by DiPRO1 and SIX1. This suggests that DiPRO1 may regulate neuronal plasticity in cancer through SIX1 targets.

In addition to SIX1, previous works dealing with the effects of myogenic regulatory factors on muscle cancer have discovered their capacity to stimulate cancer cell differentiation and deregulation of cell cycle leading to tumor growth regression (119). However, DiPRO1 seems to play a more prominent role than myogenic regulators in RMS switching them to cell death. Intriguingly, the overexpressed DiPRO1 shows co-regulation with SOX2-targets across primary recurrent or refractory childhood tumors. The role of SOX2 is related to stem neurogenesis (61) and stemness maintenance in mesenchymal (120, 121) and brain cancers (122). SOX2 could be directly regulated by SIX1 in cancer cells (122). The DiPRO1-linked gene network expectedly distinguishes RMS and Ewing sarcoma, but also medulloblastomas and neuroblastomas, suggesting that targeting DiPRO1 might have a clinical relevance for additional neuronal tumors.

Pediatric tumors overall are characterized by few specific genetic alterations (123, 124), therefore epigenetic therapy would be particularly appreciable. One recent study reported a potential efficacy of epigenetic drug combination, inhibitors of histone deacetylases (HDAC) and lysine-specific demethylase 1 (LSD1), in Ewing sarcoma xenograft models (125). Current US FDA-approved epigenetic drugs such as HDAC- and DNMT-inhibitors constitute a standard treatment in patients with leukemia, lymphoma and multiple myeloma. Despite the ever-growing progress, epigenetic medicines still need to be improved due to lack of locus specificity inducing off-target effects. The progress made in nanotechnology has allowed the development of nanomedecines based on inorganic nanoparticles for the transport and delivery of small RNA and DNA molecules, which will act with high precision and efficacy to silence target genes (126). Thus, DiPRO1-inhibiting nucleic-acid-based nanomedecines could be an attractive target for at least mesenchymal cancer treatment, contributing to the development of precision medicine. Moreover, taking into account that DiPRO1-depletion of normal cells does not induce cell death, the selective action represents a potentially less-toxic approach for treating patients. Indeed, the particular consequences of pediatric cancer treatment are related to the exposure to chemotherapy and radiation during the rapid physiologic changes that can result in specific tissue or organ damage, or alteration of growth and development. Two-thirds of survivors suffer from late effects, which could be severe or life-threatening (127). Therefore, there is a high medical need for new less toxic therapeutic approaches, particularly for pediatric cancer treatment.

## METHODS

### Antibodies

Polyclonal rabbit anti-MCM6 (Proteintech, 13347-2-AP) and polyclonal rabbit anti-TIF1B/KAP1 (Merck Millipore, Sigma-Aldrich, ABE1859), mouse monoclonal anti-ICBP90/UHRF1 (Sigma-Aldrich, MABE308), rat monoclonal anti-HA-Peroxidase high affinity (Roche, Sigma-Aldrich, 12013819001), rabbit polyclonal anti-Actin (Sigma-Aldrich, A2103), mouse monoclonal anti-FLAG® M2-Peroxidase (Sigma-Aldrich, A8592), mouse monoclonal anti-Tropomyosin Ab (Sigma-Aldrich, T2780), goat polyclonal anti-rabbit IgG–Peroxidase antibody (Sigma-Aldrich, A6154), Alexa Fluor® 488 Goat Anti-Mouse IgG antibody (Molecular Probes, Invitrogen, A-11001, A11029), Alexa Fluor® 594 Goat Anti-Rabbit IgG antibody (Molecular Probes, Invitrogen, A-11012), Fluoroshield™ with DAPI (Sigma-Aldrich, F6057), Dynabeads™ CD25 (Invitrogen, Thermo Ficher Scientific, 11157D), rat monoclonal anti-CD34 antibody (HycultBiotech, HM1015), Polink-2 Plus HRP Rat-NM Bulk kit for DAB (GBI Labs, D46-110), rabbit monoclonal anti-Ki67 (Neomarkers; LabVision, Thermo Fisher Scientific, RM-9106), rabbit polyclonal cleaved Caspase-3 (Asp175) (Cell Signaling, 9661), anti-FLAG M2 affinity gel (Sigma-Aldrich, A 2220), FLAG® Peptide (Sigma-Aldrich, F3290), Protein A agarose (Pierce, Thermo Ficher Scientific, 20333).

### Cell Culture

Immortalized human myoblasts were generated at the Institute of Myology (128) and provided by MyoBank-AFM (Paris, France). RMS JR cells (COG, RRID: CVCL_RT33) and Ewing sarcoma A673 (RRID: CVCL_0080), EW7 (RRID: CVCL_1217) cells provided by Dr C. Lanvers (University of Muenster, Germany), TE671 (RRID: CVCL_1756) provided by Dr. K. Mamchaoui (Institute of Myology, Paris, France) were STR genotyped (Mycrosynth, Switzerland). The immortalized myoblasts (Myo) were cultured in 199 medium and Dulbecco’s modified Eagle’s medium (DMEM) (Invitrogen) in a 1:4 ratio, supplemented with 20% Fetal Calf serum (FCS) (Invitrogen), 2.5 ng/ml hepatocyte growth factor (Invitrogen), 0.1 μmol/l dexamethasone (Sigma-Aldrich) and 50 μg/ml gentamycin (Invitrogen) at 37°C with 5% CO2. They were differentiated into myotubes in DMEM medium with 2% FCS and 1% penicillin/streptomycin, at 37°C with 5% CO2 for 5-8 days. The formation of multinucleated myotubes was controlled by hematoxylin staining (Sigma-Aldrich). Cancer cells were maintained in DMEM medium (Gibco, Life Technologies)/10% FBS (PAA Laboratories GmbH) with 100 UI/ml penicillin/streptomycin (Invitrogen). Cell size was measured using ImageJ software.

### Patient consent and sample collection

The cohort of this study included patients < 18 years-old at time of initial diagnosis who underwent tumor biopsy/surgery of their recurrent/refractory malignancy for WES and RNA sequencing within the MAPPYACTS (clinicaltrial.gov NCT02613962) (57). All patients and parents/legal representatives had signed informed consent for further research studies (Supplementary Tables S1-2).

### Plasmids and constructs

#### Vectors expressing the DiPRO1 ORF

Vectors expressing the ORF DiPRO1. The sequence of the full-length open reading frame (ORF) of human DiPRO1wt was optimized according to the degeneracy of the genetic code so as not to be recognized by shDiPRO1. The created cDNA (Genewiz) contained sites for the restriction enzymes XhoI and NotI for subsequent cloning into the appropriately digested retroviral vector pOZ_FHHY (129) used as a control. The resulting recombinant protein (FHHY_DiPRO1wt, pDiPRO1) contained several epitope tags and a fusion protein at the amino-terminal position: Flag polypeptide, polyhistidine motif (6xHis), hemagglutinin (HA), and the yellow fluorescent protein (YFP) (Supplementary Fig. S2A). Cell lines were infected with virus expressing pDIPRO1, and IL2Rα+/YFP+ cells were sorted using magnetic beads conjugated with antibodies against IL2Rα (dynabeads CD25, Invitrogen) and then on a FACS AriaIII cell sorter (BD).

#### Knockdown vectors targeting DiPRO1

A lentivirus-based short hairpin (sh)RNA strategy was used to repress DiPRO1 expression. The shRNAs expressed by the pKLO.1-puro vector were purchased from Sigma-Aldrich. The sequences for human shDiPRO1 were as follows:

shDiPRO1-1/sh1/shDiPRO1: CCATCATCTTTACCAATACAT

shDiPRO1-2/sh2: GCCACGTTCACCAGAAATGTT

shDiPRO1-3/sh3: CCTGAAGACAAATCCTATGAA

Myoblast cell lines were previously immortalized using a vector containing a puromycin resistance gene (128). Therefore, a second batch of shRNA constructs was created in which puromycin was replaced with blasticidin (pKLO.1-bsd). For this purpose, the cDNA of the blasticidin gene was amplified using the primers: 5’CAC ACA GGA TCC ACC GGA GCT TAC CAT GGC CAA GCC TTT GTC TC3’ and 5’CAC ACA GGT ACC GAT GCA TGG GGT CGT GCG CTC CTT TCG GTC GGG CGC TGC GGG TCG TGG GGC GGG CGT TAG CCC TCC CAC ACA TAA CCA GA3’, containing the enzymatic restriction sites *BamH*I and *Kpn*I and then inserted into the puromycin-excised-pKLO.1 vectors. Myoblasts and cancer cell lines were maintained under selection with a predetermined concentration of blasticidin S (60 µg/ml, Sigma-Aldrich). TE671 cells were also selected with puromycin (10 μg/ml). A lentivirus plasmid vector containing an shRNA insert not targeting human and mouse genes (SHC002, Sigma-Aldrich) was used as a non-target control (shCTL).

All new clones were verified by restriction enzyme digestion (*Xho*I and *Not*I) and sequencing to confirm identity (GATC Biotech).

#### Luciferase reporter vectors

The pGL3 vector-based constructs (Promega, E1751) described earlier (17) were used in these experiments. Briefly, the SV40pro (Promega, E1761) and the SV40pro+SV40e (Promega, E1741) vectors were used to evaluate promoter and enhancer activity. They contain the SV40 promoter upstream of the luciferase gene alone or with the SV40 enhancer, respectively, unlike the pGL3-basic vector without a promoter or enhancer, which was used for normalization.

### Luciferase assay

The transfection procedure and preparation of the complexes were performed using Lipofectamin 2000 reagent and 1µg of DNA according to the manufacturer’s recommended protocol (Invitrogen). The transfection experiments were performed with pGL3 luciferase reporter plasmids separatly and in co-transfection with the pDiPRO1 vector using a 1:2 ratio of DNA (µg) to Lipofectamine™ 2000 transfection reagent (µl) (Invitrogen, Life Technologies). Myoblasts and RMS cells were seeded on 24-well culture vessels at 3.8×10^4^ cells per well the day before transfection. After a 48-h incubation period, gene activity was assessed according to the supplied Luciferase Assay System Kit protocol (Promega). Luminescence detection was performed using the Microlumat LB96p device (EGG Berthold, Bad Wildbad, Germany). Protein concentration in the extracts was measured using the QuantiPro BSA Assay Kit (Sigma-Aldrich).

### Transient transfection

Cultured human TE671 cells were transfected with 1µg of DNA (pDiPRO1 or shDiPRO1) using Lipofectamine™ 2000 transfection reagent according to the manufacturer’s recommended protocol (Invitrogen, Life Technologies). Briefly, 24h before transfection, cells were trypsinized, hrvested, diluted with fresh medium, and seeded into 6-well plates at 4×10^5^ cells per well. Each sample was assayed in triplicate 48h post-transfection.

### Viral transduction

Stable overexpression or down-regulation of DiPRO1 transcripts was achieved by creating stable cell lines using lentiviral or retroviral vectors. Viral particles were obtained as described previously (129) with some modifications. To obtain transduction-competent virus, the viral vector was transfected together with pCMV and pMD2G expression plasmids for gag-pol and env. The HEK293T (RRID: CVCL_0063) cells were transfected using the calcium phosphate transfection method (CAPHOS, Sigma Aldrich). Selection was started 4-5 days post-infection using either magnetic sorting (pOZ), antibiotic selection (pOZ, pLKO.1-puro/blast), or fluorescence sorting (pOZ). For antibiotic selection, the lethal concentration of the antibiotic was first tested on parental nontransduced cells and used as a reference to select stable cell lines (10 μg/mL puromycin or 60 μg/mL blasticidin).

### Assessment of cell differentiation and cell dimensions

Differentiation was assessed morphologically based on May-Grünwald–Giemsa staining. To this aim, cells were cytospun onto slides, fixed with absolute methanol for 15 min at room temperature and then transferred to a staining tank containing May-Grünwald dye diluted in an equal amount of water for 5 minutes and then to the Giemsa solution (Unité de cytopathologie, Gustave-Roussy). The extent of myogenic differentiation was determined by the number of multinucleated myotubes formed and the expression level of myogenic regulatory factors (tropomyosin). Cells containing more than three nuclei were concidered as multinucleated myotubes. and then at least 20 cells for each experimental condition were analyzed on an Olympus IX83 inverted microscope equipped with an ORCA-Flash 4.0 digital CMOS camera (HAMAMATSU). Alternatively, the glass slides were scanned and real images were taken using an EVOS M7000 Imaging System (Thermo Fisher Scientific). The images were visualized using Olympus OliVIA software. Cell size and nuclear-cytoplasmic ratio (N/C) (42) was scaled and evaluated using ImageJ.

### Cell viability and proliferation assays

Cells were subcultured in multi-well coated cell culture plates and allowed to reach ∼95% confluence within 72 h. Cell viability was assessed using Cell Counting Kit-8 (Sigma-Aldrich), according to the supplier’s recommendations. Briefly, WST-8, a tetrazolium salt, is reduced by cellular NAD+- and NADP+-dependent dehydrogenases to an orange formazan product soluble in tissue culture medium. The amount of formazan formed is directly proportional to the number of living cells. The number of cells was determined using cells at different dilutions: from 0 to 25,000 cells/well/100 μL. Absorbance was measured at 460 nm using a plate reader 1h after seeding. The resulting titration curve was used to estimate the number of viable cells for proliferation assay. Cells were then seeded at 1.0×10^4^, 1.5×10^4^ and 2.5×10^4^ cells/well and absorbance was measured after 72h of proliferation. Data are expressed as the average percentage of three dilutions of cells after 72h relative to the initial number of cells (set at 100 %).

### Immunofluorescence microscopy

Cells were grown on 2-well chamber slides (Nunc, Lab-Tek) and fixed in 4% formaldehyde, permeabilized with 0.2% Triton, and blocked in PBS containing 3% bovine serum albumin (Sigma) for 1 hour at room temperature. The cells were then incubated with anti-Tropomyosin monoclonal Ab (1/50, Sigma-Aldrich) and anti-actin Ab (1/100, Sigma-Aldrich) for 1 hour. Signals were detected with the appropriate Alexa 488 or Alexa 594-conjugated secondary antibodies (1/100, Molecular Probes Invitrogen) for 1 hour at room temperature. For nuclear counterstaining, cells were mounted with Fluoroshield containing DAPI (1/3, Sigma-Aldrich). Images were obtained using an Olympus IX83 inverted microscope equipped with an ORCA-Flash 4.0 Digital CMOS camera (HAMAMATSU).

### Gene expression analysis

#### RNA isolation and RT-qPCR

Total RNA was extracted using the RNeasy Mini Kit (Qiagen) for cell culture samples. cDNA was synthesized from 500 ng of total RNA using the RevertAid H Minus First Strand cDNA Synthesis Kit (Fermentas, Thermo Fisher Scientific) for RT-PCR analysis according to the manufacturer’s instructions. n the StepOnePlus Real-Time PCR Detection System (Applied Biosystems) using FastStart Universal SYBR Green Master Mix (Rox) (Roche Life Science). Relative expression was determined by the 2^-ΔΔCt^ method (130) using GAPDH as an internal control. The oligonucleotide sequences used in this study are listed in Supplementary Table S3.

#### Microarray assay

For microarray analysis, RNA quality was assessed using a Bioanalyzer (Agilent) and then quantified using a Biospecnano instrument (Shimadzu, Kyoto, Japan). Gene expression analysis was performed using an Agilent® SurePrint G3 Human GE 8×60K Microarray (Agilent Technologies, Santa Clara, CA, USA) using an Agilent Single Color Labeling Kit (Low Input Quick Amp Labeling Kit 034949) adapted for small amounts of total RNA (100 ng total RNA per reaction). Hybridization was performed according to the manufacturer’s instructions. Microarray images were analyzed using Feature Extraction software version (10.7.3.1) from Agilent technologies. Defaults settings were used. Raw data were processed using R with LIMMA, the R package from the Bioconductor project, and processed as follows: the gMedianSignal and rMedianSignal data were imported, the control probes were systematically removed, and flagged probes (gIsSaturated, gIsFeatpopnOL, gIsFeatNonUnifOL, rIsSaturated, rIsFeatpopnOL, rIsFeatNonUnifOL) were set to NA. To obtain a single value for each transcript, the average value of the total data of each replicated probe was taken. The missing values were calculated using the KNN algorithm from the ’impute’ package from the R Bioconductor. The normalized data were then analyzed. To estimate the differentially expressed genes between the two groups, we began by fitting a linear model to the data. We then used an empirical Bayesian method to moderate the standard errors of the estimated logarithmic changes. The top ranked genes were selected according to the following criteria: absolute fold change (FC) ≥ 1.4 and ≤-1.4 and p-value < 0.05.

#### RNA-seq

RNA-seq libraries from the MAPPYACTS study sequencing data were aligned with the GRCh38 version of the human transcriptome from the GENCODE project using Salmon v0.9 (131) and converted from transcripts to gene expression using the Tximport R package. RNA-seq analysis data of 1375 cancer cell lines were extracted from the DepMap Broad Institute portal. The data were then computed using DEBrowser (132) and the DESeq2 R package (TMM normalization, parametric fit, LRT test). Coexpression analysis was performed using the network_plot R function.

### Flow cytometry analysis

Cells were washed with cold PBS, fixed in cold 70% ethanol for at least 30 min. The supernatants were washed twice in PBS. The cells were then treated with RNase (Macherey-Nagel, Thermo Fisher Scientific), stained with propidium iodide (PI) (Sigma-Aldrich), DNA content and percentage of viable and dead cells in each phase of the cell cycle were assessed using a C6 Accuri flow cytometer (BD Biosciences) and FlowJo-V10 software (FlowJo, LLC).

### Immunoaffinity purification

DiPRO1 ORF was cloned into the pFHHY-expression vector, and a stable cell line overexpressing the DiPRO1wt protein (pDiPRO1) was generated. Immuneaffinity purification was based on the protocol proposed in (129) (133) and optimized by our team. Approximately 10×10^6^ RMS cells were lysed in CSK buffer (1:5 package V) containing 10 mM Tris-HCl, pH 7.5 100 mM NaCl, 300mM sucrose, 3mM MgCl_2_, 1 mM EGTA, 1 mM NaF, 1 mM Na3VO4, 0.5% Triton X-100 and 2 mM PMSF, Complete™ EDTA-free Protease Inhibitor Cocktail (Sigma Aldrich), and 50 μL were taken for the input lanes. After centrifugation for 10 min at 4000 rpm, the supernatant (cytosolic fraction) and pellet (nuclear fraction) were collected. Nuclei were then resuspended in nuclear incubation buffer (150 mM Hepes, pH 7.9; 1.5 mM MgCl2; 150 mM KOAc; 10% glycerol; and protease inhibitors) and digested with 0.15 unit/μl benzonase (Sigma Aldrich) according to Aigun et al. (134). The samples were cleared by centrifugation at 20,000 × *g* for 20 min, and proteins were collected from the nuclear extract. The identical procedure was used to purify isogenic cells lacking bait protein expression (null control).

Nuclear extracts were pre-cleared with an inert resin Protein A Agarose (Thermo Fisher Pierce) for 2h at 4°C with rotation, then separately applied to ANTI-FLAG M2 Affinity beads (Sigma-Aldrich) (≈ 25 μl of packed beads) and incubated for 2 h at 4°C. After binding the protein complexes, the beads were washed extensively with wash buffer [1mL 2M Tris HCL pH7.4, 25mL Glycerol 99.5%, 150µL 1M MgCl2, 40µL 0.5M EDTA pH8, 9.99mL 3M KCL, 200µL 100mM PMSF, 70µL 14.3M bME, and protease inhibitors]. Finally, the purified protein complexes were eluted with FLAG elution buffer [1mL 2M Tris HCL pH7.4, 25mL Glycerol 99.5%, 150µL 1M MgCl2, 40µL 0.5M EDTA pH8, 9.99mL 3M NaCL, 400 μg/ml 3× FLAG peptide, 200µL 100mM PMSF and protease inhibitors] or/and with LDS loading buffer. Eluates were resolved in 4–12% bis-Tris gradient PAGE (NuPAGE, Life Technologies) and Western blotting was perfomed using antibodies raised to HA-epitope (Sigma Alrich). When the protein bands were excised for mass spectrometry, the gels were stained with InstantBlue (Expedeon).

### Sample preparation and proteomic analysis by MS/MS

#### Sample preparation

The Co-IP-derived gels were combined and underwent in-gel digestion as described previously (135). Briefly, the gel slices were washed with 100 μl of 1:1 acetonitrile (ACN):50mM Ambic followed twice by 100 μl of 100% ACN, then reconstituted with 100 μl 10 mM DTT and alkylated with 100µl 55mM Iodoacetamide. Final dehydration was performed using 100 μl of 100% acetonitrile. Each wash was carried out for 20 min at 37°C with shaking at 1400 rpm. The gel slices were dried in a speed-vac for 5 min. Gel slices were pre-incubated with 20 μl of pre-activated trypsin (Promega France, Charbonnières) at room temperature for 15-20 min, 50 mM ammonium bicarbonate was added to coat the gel, and incubated at 37°C for 16 h. Peptide-containing supernatants were collected, and the gel slices were washed twice with 50µl of 0.1% formic acid in 70% ACN for 15min at 37°C. The collected supernatants were dried on a high-speed fan until completely dry, then resuspended in 10 μl of water with 0.1% formic acid (v/v) for mass spectrometric analysis.

#### MS/MS analysis

Protein identification were performed using an Orbitrap Q-Exactive mass spectrometer (Proteomic Platform, Gustave Roussy) supplied with an Easy-nLC-100 chromatography system (Thermo Scientific) coupled to an Orbitrap Q-Exactive analyser (Thermo Scientific) and an Easy-spray column source for samples ionization (Thermo Scientific). The gradient used for protein identification and/or MS2 targeting was as follows: 0 to 120 min: 5% to 35% solution B; 120 to 140 min: 35 to 50% solution B; 140 to 145 min: 50 to 90% solution B; 145 to 150 min: 90% solution B, where solution A: 100% water with 0.1% formic acid and solution B: 100% acetonitrile and 0.1% formic acid. The MaxQuant package was used to identify proteins using the human genome FASTA database. The software was adjusted to a confidence level of 10ppm for MS and 0.02 Da for MS/MS. The p value used for peptide confidence identification was p≤0.01. The obtained proteins were then analyzed using Perseus tools (136).

#### Mass spectrometry data processing

Mass spectrometry data processing. The data obtained from the mass spectrometry analysis were then filtered according to the following parameters: (a) at least 2 unique peptides must match the sequence of the candidate protein, and (b) the proteins must show a significant (p ≤ 0.05) increase in LFQ intensity in the DiPRO1 fraction compared with the control fraction. Common false positives, such as keratins, keratin-associated proteins, and serum albumin were excluded from the analysis. In addition, the CRAPome database [http://www.crapome.org, (137)] was employed to exclude the most likely false positive hits.

#### Validation of DiPRO1-protein binding

The DiPRO1-interactions were confirmed with co-immunoprecipitation (Co-IP) using specific antibodies and Protein A Agarose (Thermo Fisher Pierce). Nuclear extracts were isolated as described in the Immunoaffinity purification section. Specific antibodies (5 µg) were separately incubated overnight at 4 °C with 500 µg of protein with constant rotation. Protein A agarose beads were added (100 μl of 50% slurry in lysis buffer) and incubation was performed for 2 h at 4 °C with rotation. The mixture was washed three times and centrifuged for 3 min at 500g at 4 °C. Precipitated proteins were eluted by boiling in NuPAGE LDS sample buffer and analyzed by Western blotting using appropriate antibodies. Interaction with DiPRO1 was included as a positive control. To this end, DiPRO1 was co-immunoprecipitated from cell lines expressing DiPRO1 using the anti-FLAG antibody. Protein complexes were analyzed by Western blotting using HRP-conjugated anti-HA antibodies.

### Protein isolation and immunoblotting

Sample pellets were diluted in 150-300 μl of 1× phosphate-buffered saline with protease inhibitor cocktail (Sigma-Aldrich) and sonicated. NuPAGE® LDS sample buffer (Life Technologies) and 5 mM DTT were added and the samples were incubated for 30 minutes at 50°C. Western blotting was performed according to a standard procedure using pre-cast gel cassettes (Life Technologies) and NuPAGE® MOPS SDS run buffer (Thermo Fisher Scientific). The Pierce G2 Fast Blotter device (Thermo Fisher Scientific) was exploited for semi-dry transfer of protein gels. SeeBlue Plus2 protein standard (Life Technologies) was used to detect the size of the protein being analyzed. The detection solution was HRP Immobilon Western Substrate (Merck Millipore). The analysis was performed using ANTI-FLAG® M2-Peroxidase (Sigma-Aldrich, 1/5,000), anti-HA-peroxidase (Roche, Sigma-Aldrich, 1/1,000), anti-Actin (Sigma-Aldrich, 1/3,000) and anti-rabbit secondary antibodies (Sigma-Aldrich) in 1/1000 dilution. Fold change was calculated after quantification using ImageJ software.

### ChIP-seq

To associate the binding site information with gene expression, we performed the analyses as outlined below. First, the ChIP-seq processed datasets of DiPRO1/ZNF555 from reference (10) (NCBI’s GEO accession number GSM2466593) were genome-wide annotated using the HOMER package (138) and the PAVIS tool (139). The target region was located 10 Kb upstream of the TSS and 5 Kb downstream of the TES. The annotation of repeats was realized using the EaSeq ‘Colocalization” tool and the UCSC “rmsk” track. In the ChIP experiment, 293T cells were induced to express a cDNA encoding human DiPRO1 by lentiviral transduction. For expression-linkage correlation analysis, our series of gene expression experiments (p < 0.05) reported here were used in combination with ChIP-seq to define a set of genes that are enriched in direct transcriptional targets. Motif enrichment and similarity analysis were performed using Homer’s tool and the MEME suite tool (140): MAST, FIMO (141), Tomtom (142). Cis-regulatory regions were determined using GREAT software (default settings for cis-regions: 5.0 kb constitutive upstream and 1.0 kb downstream of TSS regions and up to 1Mb of extension) (143). The ChIP-seq data were remapped to human RefSeq hg38 using the NCBI genome “Remap” tool. Transcription factor binding regions and chromatin state were annotated using the ENCODE TxnFactrChIPE3 and BroadChromHMM tracks (37, 38).

### MIRA-seq

#### DNA extraction and library preparation

Cells were washed in PBS and mixed with 500 µl of ice-cold MES lysis buffer (144), incubated for 5 minutes and centrifuged at 10,000 rpm for 5 minutes. The supernatant was collected and then diluted in 300 µl of LDS extraction buffer (10% glycerol, 0.14 M Tris-Base, 0.14M Tris-HCl, 2% LDS, 500nM EDTA. The samples were briefly sonicated to reduce viscosity and then centrifuged. The resulting supernatant was mixed with an equal volume of phenol:chloroform:isoamyl alcohol (25:24:1 [vol/vol]). The aqueous phase was retained by centrifugation, and then the dissolved DNA was precipitated with 0.1 volume of 2.5 M NaCl and a 2.5-fold volume of ice-cold ethanol. The samples were then centrifuged at 15,000×g for 15 min at 4°C. Finally, the DNA pellet was washed with precooled 70% ethanol and resuspended in 50 μL of Milli-Q water. The concentration of extracted DNA was measured using the Qubit instrument and the Qubit dsDNA HS Assay Kit (Thermo Fisher Scientific). After extraction, genomic DNA was sonicated (300-700 bp size) and MIRA pull-down was performed using the Methyl Collector Ultra Kit (Active Motif), which uses a MBD2b/MBD3L1 protein complex, according to the manufacturer’s instructions starting with 1 µg of DNA and using high salt binding conditions. Extracted genomic DNA (gDNA) was sonicated in a Covaris S220 (LGC Genomics GmbH) to an average fragment size of 180 bp. Fragment length distribution was assessed by microelectrophoresis using the Bioanalyzer 2100 (Agilent). Enriched DNA and input DNA fragments were end-repaired, extended with an ’A’ base at the 3′ end, ligated with pair-indexed adapters (NEXTflex, Bioo Scientific) using the Bravo platform (Agilent), and PCR amplified for 10 cycles. Final libraries were purified with AMPure XP beads (Beckman Coulter), pooled in equal concentrations, and subjected to pair-end sequencing (100 cycles: 2×50) on the Novaseq-6000 sequencer (Illumina) at the Genomic platform (Gustave Roussy, Villejuif, France).

#### Bioinformatics analysis

Using the Galaxy platform (145), reads were groomed and cleaned by clipping (ClipAdapter) and trimming (TrimGalor) and the quality of NGC data was evaluated using the FastQC tool. GC content per sequence was distributed normally for input samples and showed an approximately 6-10% higher shift for MIRA-seq reads. The Bowtie2 algorithm was used to map the pared end reads against the hg38 genome index version 2013. The resulting BAM formatted files were filtered using bitwise flag-based alignment records to obtain the output of uniquely mapped reads, which were used for downstream analysis. Bedtools were used to convert BAM to BED. The following steps of the analysis were performed using the free interactive software EaSeq (146). Easeq browser was used for tracking and visualization of genomic regions. Peak searching was performed with the default procedure, adaptive local thresholding (ALT). The settings were window size 200 bp, p-value ≤ 10^-5^, FDR 10^-5^, and Log2FC 2. Peaks were merged within a 500 bp size. CpG sites were downloaded from http://methylqa.sourceforge.net/. The human genome (version hg38) and repeat-masked CGI (hk-CGI) and unmasked CGI tracks were downloaded from the UCSC genome database (147). The rep-CGI regions correspond to the unmasked CGI excluding hk-CGI. CGI meets the following criteria: GC content of 50% or more, length greater than 200 bp, ratio greater than 0.6 between the observed number of CG dinucleotides and the expected number based on the number of Gs and Cs in the segment. The genomic distribution of CGIs was analyzed by the annotation tool with the hg38 genome, and regions from 501 to 10,000 bp upstream of transcription start sites (TSSs) were considered as promoter regions, 500 bp upstream and downstream of TSSs were considered as TSS regions. CGI repeats were annotated using the “Colocalization” tool with RepeatMasker track from UCSC Genome Browser.

#### *In vivo* study design

All procedures on animals were approved by the CEEA26 Ethics Committee, and the Ministry of Agriculture (approval number: APAFIS#8550-201701160922201) and were performed within the guidelines of humane care of laboratory animals established by the European Community (Directive 2010/63/UE). The antitumor activity of DiPRO1 inhibition was evaluated using PEI-nanocomposites containing siRNA oligos or a vector expressing a DiPRO1-targeting shRNA compared with a non-targeting siRNA. Nucleic acids (10 μg) were complexed with a linear polyethylenimine transfection agent (*in vivo*-jetPEI ®; Polyplus Transfection) at an N/P ratio of 8 (N/P ratio is the number of nitrogen residues in jetPEI ® per nucleic acid phosphate) and resuspended in 50 μL of 5% (wt/vol) glucose. The shDiPRO1 conformed to shDiPRO1-1 (see section Plasmids and Constructs). The modified siRNA-based sequences were as follows:

siDiPRO1:
Sense: 5’-GCCAUCAUCUUUACCAAUAdTdTdTdTdT-3’
Antisense: 5’-UAUUGGUAAAGAUGAUGGCdAdAdAdAdA-3’
Scramble control siCtl (148)
Sense: 5’-AUGUCUACUGGCAGUCCUGdTdTdTdTdT-3’
Antisense: 5’-CAGGACUGCCAGUAGACAUdAdAdAdAdA-3’
siDiPRO1_Cy5
Sense 5’-[CY5]GCCAUCAUCUUUACCAAUAdTdTdTdTdT-3’
Antisense: 5’-UAUUGGUAAAGAUGAUGGCdAdAdAdAdA-3’
d is deoxyribonucleotide

For subcutaneous (s.c.) xenograft models, 6×10^6^ Ewing sarcoma (A673) cells were injected into the right flank of six-week-old females (Hsd:Athymic Nude-Foxn1^nu^, Envigo, 6903F). Animals with established s.c. tumors were randomly assigned to treatment groups (n=7 or 5) receiving either nanocomposites of (i) shDiPRO1/jetPEI®, (ii) shDiPRO1/jetPEI® or (iii) scramble siRNA controls (Ctl). In accordance with the “3 R’s” of animal research, we did not use the second scramble control (shRNA), considering the lack of effect *in vitro* as identical to scrambled siRNA. The nanocomposites were administered intratumorally in glucose solution at 0.5 or 1.0 mg/kg/injection twice or three-times per week. The mice were monitored regularly for changes in tumor size and weight. Animals were sacrificed according to the animal ethic endpoints or at the end of the experiments. Satellite mice were treated equally to the treatment groups and sacrificed for histochemical and tumor cell uptake analysis using fluorescent nanocomposites coupled to Cy5. Fluorescence emission was monitored at 24 and 72 h using the 3D scanner of the IVIS® Spectrum *in vivo* imaging system (PerkinElmer) and analyzed by Living Image Version 4.5.2 software after a single administration at 0.5 mg/kg/injection. The length and width of the tumor were measured with calipers, and the Antitumor activity was assessed on the basis of tumor growth delay (TGD) in median time for tumor volumes to be doubled (149), and tumor growth inhibition at day 10 (TGId10, %) according to (150). Toxicity was defined as body weight loss >20%, or mortality. Data analysis was performed using GraphPad Prism and TumGrowth (151) software. Statistical tests and the number of biological replicates (n) for each experiment are indicated in the figure legends.

### *In vivo* tumor cell internalization of amino-acids/PEI particles

*In vivo* tumor cell uptake was measured in mice with siDiPRO1 coupled to the fluorescent red dye Cy^5^ (Sigma-Aldrich) delivered by intratumoral injection at 0.5 mg/kg. Fluorescent images were followed 24 and 72 h post-injection with the 3D scanner of the *in vivo* imaging system (IVIS® 200 Spectrum CT, PerkinElmer) and analyzed by the Living Image Software Version 4.5.2.18424. In addition, tumors were extracted and fluorescence monitored *ex vivo*.

### Immunohistochemical analysis of xenografts

Tumors were fixed in 4% paraformaldehyde, embedded in paraffin, and sections (4μm) were stained with hematoxylin eosin and safranin (HES). For detection of *in situ* cell death (Roche), sections were processed for proteolytic digestion with proteinase K (Roche) and incubated with TUNEL solution. Staining was visualized using the permanent red kit (Dako) and examined using a Zeiss Axiophot microscope. For immunohistochemistry, sections were incubated after heat-induced antigen retrieval with rat anti-mouse CD34 antibody (1/20) (Hycult technologies), rabbit monoclonal anti-Ki67 (1/200) (Neomarkers; LabVision), anti-cleaved caspase 3 (1/100) (Cell Signaling), visualized using the polink anti rat kit (GBI Labs) and the peroxidase/diaminobenzidine Rabbit PowerVision kit (ImmunoVision Technologies), respectively. Slides were acquired with a Virtual Slides microscope VS120-SL (Olympus, Tokyo, Japan), 20X air objective (0.75 NA) and visualized using OLYMPUS OlyVIA software. TUNEL-stained cells were detected using the Definiens Tissue Studio software nuclei detection algorithm (Definiens AG, Munich, Germany). The analysis was performed on manually selected regions of interest by a qualified pathologist. Briefly, stained nuclei were automatically detected according to their spectral staining properties in the regions of interest. Thresholds on the area of the nuclei, set at 10μm² and 60μm², were used to discard objects that were too small or too large from the analysis. The algorithm exports for each region of interest the number of stained cells and the total area in µm². The quantitative analysis represents the % of positive cells in the tumor tissue sections compared to the total number of tumor cells. Five to eleven fields were chosen for counting. ImmunoRatio ImageJ plagin was used to quantify Caspase 3, Ki-67 and CD34 positive cells. The average count in each region was used for statistical analysis by t-test. Necrotic fields were excluded. Results are means ± SEM of at least three independent experiments. Statistic tests and number of biological replicates (n) per each experiment are outlined in figure legends.

### Analysis of somatic mutations and copy number variations

MAPPYACTS samples were sequenced using the Illumina NextSeq 500 or Hiseq 2000/2500/4000 platforms [UMO1] to produce library of 75-bp or 100-bp reads. Three different kits were used to perform the Exome capture: SureSelect XT human All exon CRE version 1 or 2 and Twist Human Core Exome Enrichment System. Sequencing libraries quality was estimated with fastqc and fastqscreen. Reads were mapped with BWA (v0.7.17 with parameters: -M -A 2 - E) onto the Human reference genome assembly hg19/GRCh37 120. SNVs (single nucleotide variations) and small indels were called using GATK3 121 (Indel Realigner, Base Recalibrator), samtools (fixmate, markdup, mpileup) and Varscan (v2.3.9) from paired normal/tumors bam files. Variant annotation and functional prediction was performed with ANNOVAR 122 using public database releases on 2019.11.07 from 1000 genomes project, Exome Aggregation Consortium, NHLBI-ESP project, Kaviar. Somatic mutations were filtered according to their enrichment in the tumor samples compared to the paired normal samples using a Fisher test and p-value<0.001. To estimate mutational spectrum, we excluded questionable somatic variants observed in less than 3 reads, or with an allele frequency lower than 0.05, or described in 1000 genomes and EXAC databases with a frequency higher than 0.05%. Non exonic variant, not covered by the Exome capture, were also excluded. Additionally, we performed the allele-specific copy number profiling using Sequenza algorithm (152). Focusing on the genomic region of DiPRO1/ZNF555 in our cohort study, segments with start or end in, or start before and end after, were considered to be related to the ZNF555 gene. CNV (copy number variation) was defined as gain: CN > ploidy; loss: CN < ploidy; LOH (Loss Of Heterozygosity): deletion of one allele; LOH copy neutral (cn): deletion of one allele, with a gain of the second allele and ultimately a copy number equal to the sample ploidy; Homozygous deletion: no DNA copy. Amplification was defined starting from 7 CN for a ploidy of 2. For 3≤ploidy≥ 9, the DNA copy cutoff was set according to the expression: ploidy x 2 + 1; and for 10 ≥ ploidy, the DNA copy cutoff was set according to: ploidy + 11. For the correlation between gene expression and copy number, gene-wise CNV was regarded as -3 = homozygous deletion; -2 = hemizygous loss LOH; 1 = hemizygous loss without allele deletion; 0 = neutral/cn; 1 = gain without allele deletion; 2 = gain LOH; 3 = amplification. Z scores of gene expression levels were computed using tumors samples that are diploid for the corresponding gene. For each sample, the Z score = (x-u)/o (65), where u, o represent the mean and standard deviation of the DiPRO1 gene expression among diploid samples for the DiPRO1 gene, respectively; x represents the DiPRO1 gene expression in a sample with CNV. As illustrated in Supplementary Fig. S5A, the variation in DiPRO1 expression level is moderate and thus, differentially expressed DiPRO1 was considered upregulated when Z scores ˃ 0.5 or downregulated when Z scores <− 0.5. Spearman’s correlation coefficient (r) test and nonlinear regression analysis were implemented to define the relationship between differential DiPRO1 gene expression and CNV using GraphPad Prsim.

### Functional analysis

Gene ontology term enrichment was analyzed using GOrilla or Cytoscape software with the ClueGO and the CluePedia plug-ins (apps.cytoscape.org/apps/cluego) (153) to decipher gene ontology annotation networks and functionally clustered pathways. Benjamini-corrected term values of p < 0.05 were considered statistically significant. The two-tailed hypergeometric test and Bonferroni step down correction method were used. Group values of p < 0.05 were considered statistically significant. The parameters were as follows: Kappa score = 0.4, H. sapiens [9606], genes in GO_BiologicalProcess-EBI-QuickGO-GOA. GO annotation groups “GO_Muscle” and “GO_CellCycle” were downloaded from the Gene Ontology annotation database. Regulator prediction was performed using the iRegulon V1.3 (60). Positional gene enrichment analysis was performed using the PGE tool (80). Molecular Signatures DataBase (MSigDB) and Gene set Enrichment (GSEA) software were employed to analyze the gene set according to their hallmarks and target genes (154). The Venn diagrams were made using Venny 2.1 (<u>bioinfogp.cnb.csic.es/tools/venny/</u>) and Jvenn tools (155).

## Supporting information

Duppl Fig. S1-7 and Tables S1-3

## ACKNOWLEDGEMENTS

We are grateful to Prof. Pascal MAIRE (Inserm INSERM U1016, Paris, France) for the critical reading of the manuscript. We are grateful to Prof. Valisly OGRYZKO for his contribution and helpful advice on the development of the experimental strategy. We thank Dr. Andrey MAKSIMENKO (Harrison School of Pharmacy, Auburn University, USA) and Dr. Estelle DAUDIGEOS (Gustave Roussy, Villejuif, France) for their expert assistance in the design of the nanomedicine and *in vivo* experiments. We also thank the proteomics facilities for the qualified validation of the proteomic data (proteomic platform of the Institut Gustave Roussy, Villejuif; 3P5 of the Université Paris-Descartes) and François GILLETTE (Hays) for the computer assistance This study used patient data collected as part of the MAPPYACTS trial supported by the PHRC-K14-175 grant from the Institut National du Cancer, the MAPY201501241 grant from the ARC Foundation and the Association Imagine pour Margo. We thank the Encode Consortium and the Encode production laboratories that generated the data sets. We thank the MyoBank AFM-Institut de Myologie (Paris, BB-0033-00012), Dr. Vincent MOULY and Dr. Kamel MAMCHAOUI (Sorbonne University, Inserm, Institut de Myologie, U974, Centre de Recherche en Myologie, Paris) for providing myoblasts. We would like to express our gratitude to Laurent BERNARD and Mathieu POIROT (DRH, CNRS Délégation Paris-Villejuif) for their administrative support and to Corinne GAULTON for the copy editing.

## DEDICATION

Dedicated to the talented scientists, Dr. Andrey Maksimenko, Harrison School of Pharmacy, Auburn University, USA, and Dr. Vaslily Ogryzko, INSERM, CNRS, Gustave Roussy, France, who contributed significantly to this work and passed away unexpectedly.

