## Supplementary material for "DiPRO1 dependent transcriptional and epigenetic regulation distinctly controls the fate of muscle and mesenchymal cancer cells": Duppl Fig. S1-7 and Tables S1-3

### Supplementary Figure S1

**A** DiPRO1/ZNF555 protein

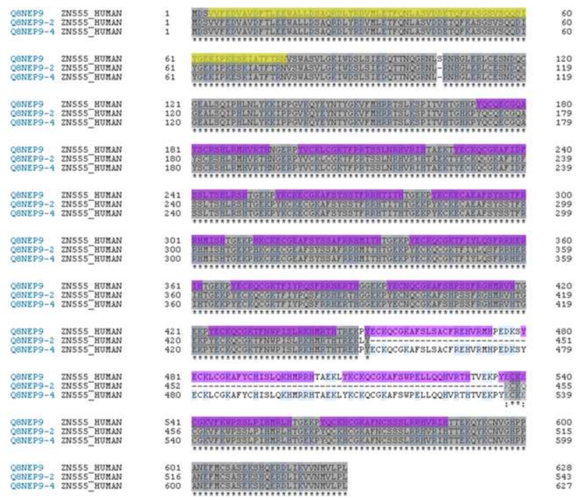

- KRAB A
- Zn-fingers (12-15)
- Identical for 3 isoforms

Predicted size

Isoforme 1 : 73.1 kDa (Q8NEP9-1) 628AA

Isoforme 2 : 63.0 kDa (Q8NEP9-2) 543AA

Isoforme 3 : 73.0 kDa (Q8NEP9-4) 627AA

**C**

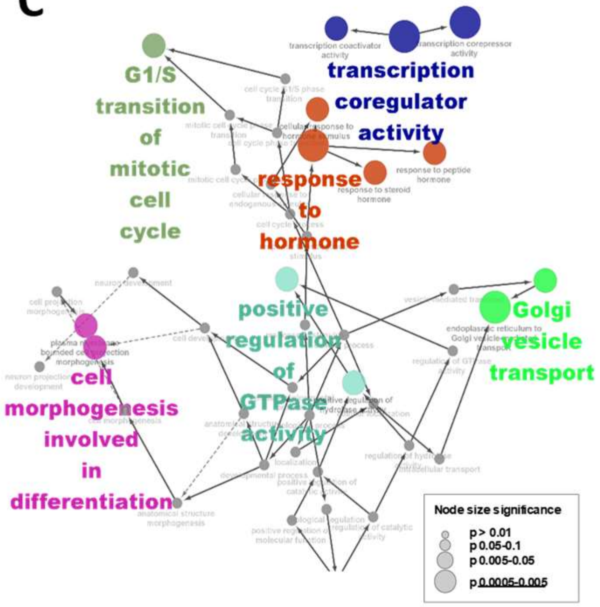


**B**

| Chromatin annotation | % |
|---|---|
| Transcriptional elongation | 6.34 |
| Transcriptional transition | 2.38 |
| Weak Transcription | 15.45 |
| Weak Promoter | 0.59 |
| Active Promoter | 0.40 |
| Strong Enhancer | 2.18 |
| Weak Enhancer | 3.37 |
| Polycomb -repressed | 3.17 |
| Heterochrom low signal | 60.00 |
| Repetitive/Copy Number Variation | 0.79 |
| Poised Promoter | 0.40 |
| Insulator | 0.40 |
| NA | 4.55 |

**D**

| Rank | Motif | P-value | % of targets | % of BG | Best match |
|---|---|---|---|---|---|
| 1 | 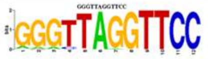 | 1e-204 | 16.04 | 0.01 | **Six1** (GSE20150) |
| 2 | 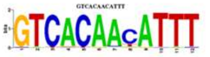 | 1e-84 | 6.93 | 0.05 | **Foxj1** (PB0120.1) |
| 3 | 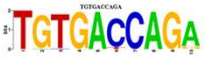 | 1e-76 | 8.71 | 0.17 | **Nr2f2** (PB0153.1) |
| 4 | 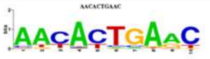 | 1e-69 | 12.28 | 0.85 | **Nkx3.1** (GSE28264) |
| 5 | 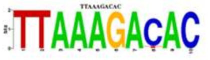 | 1e-62 | 5.35 | 0.16 | **Hox9** (PH0073.1) |
| 6 | 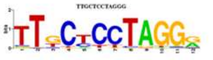 | 1e-58 | 7.92 | 0.03 | **MZF1** (MA0057.1) |
| 7 | 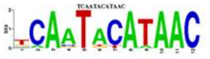 | 1e-49 | 6.34 | 0.01 | **HNF1B** (MA0153.1) |
| 8 | 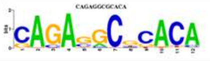 | 1e-46 | 37.82 | 8.85 | **Gm397/ZSCAN4** (PB0130.1) |

**E**

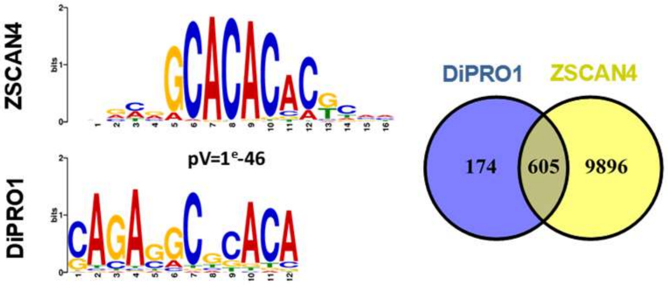


**F**

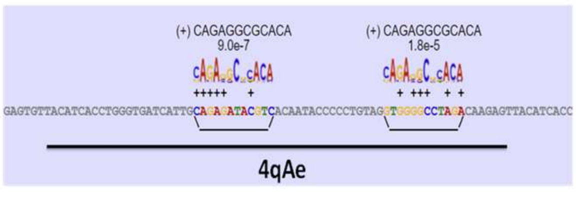

#### Supplementary Fig. S1. Supplementary Figure S1. DiPRO1 direct targets and their functions.

**A,** Amino-acid sequence alignment of three DiPRO1/ZNF555 isoforms. Residues shaded in blue correspond to charged amino acids. **B-F,** ChIP-seq data for DiPRO1 (GSM2466593) performed in HEK 293T cells. **B,** Chromatin annotation of DiPRO1 binding regions. **C,** A network of functional groups of enriched categories was created for the DiPRO1 target genes. GO biological process (BP) groups of enriched terms are represented by colored nodes, and node size reflects the significance of term enrichment, p-value group corrected with Bonferroni stepwise reduction, kappa score = 0.4. Gray nodes correspond to parent nodes connected by flashes to root nodes. **D**, Top 8 *de novo* motifs for DiPRO1 and the best matches. Motives from the ChIP-seq experimental data were prioritized. **E**, The most abundant DiPRO1 binding motif 8 and its best match ZSCAN4 (left). The motif alignment was shown for the ZSCAN4 human PB0130_1/Jaspar using the Tomtom tool. Gene overlap between the *cis*-regulatory regions of DiPRO1 and ZSCAN4 (right). **F**, The DiPRO1 motif 8 matches the beta satellite DNA repeat of the 4qAe sequence.

### Supplementary Figure S2

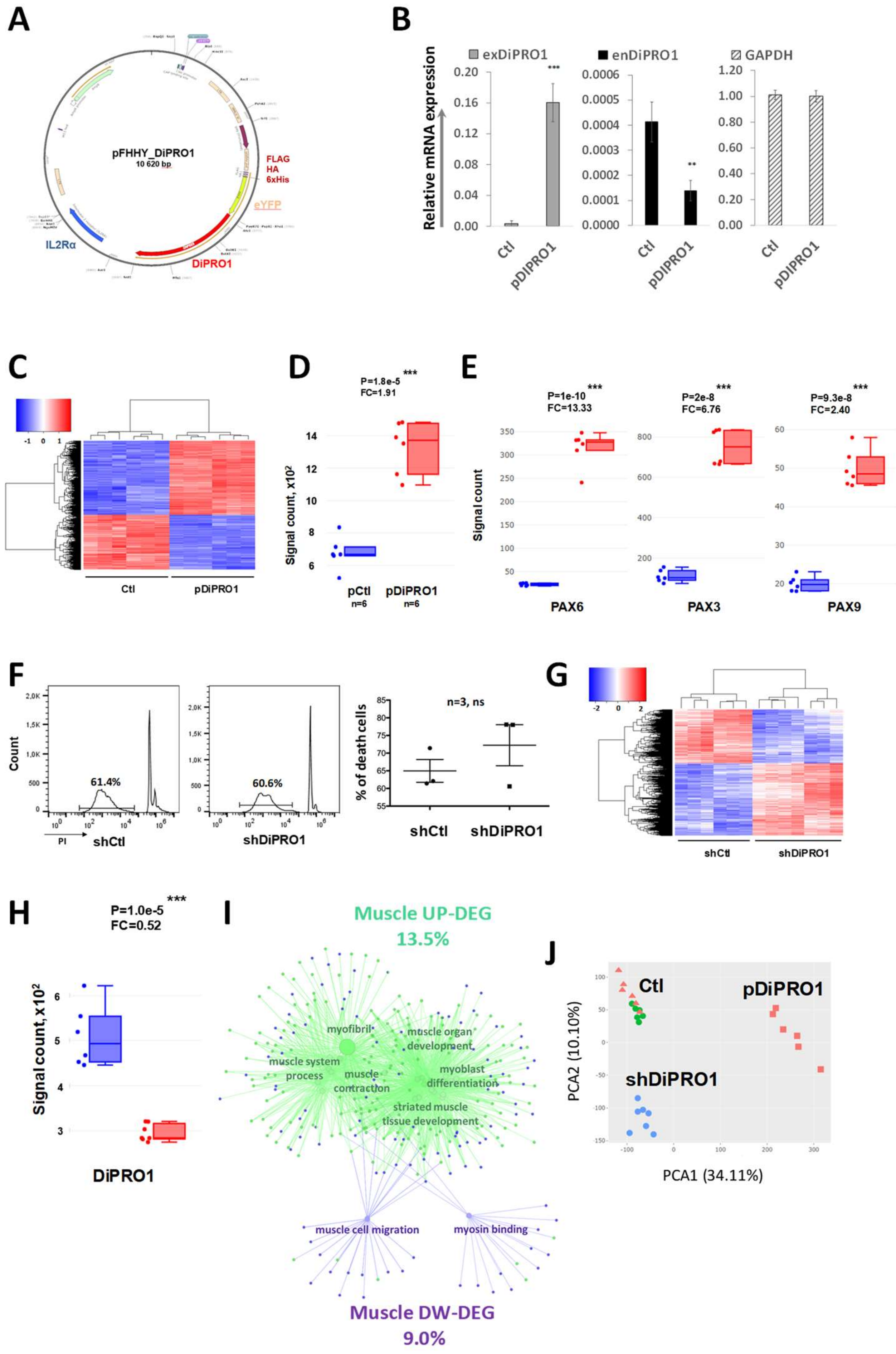

#### Supplementary Figure S2. DiPRO1 regulates myogenic genes of human myoblasts.

**A,** Map of the pDiPRO1 retroviral vector showing its major components. The DiPRO1 ORF was inserted into the pOZ vector backbone between the XhoI and NotI sites, with the epitope tags FHHY: FLAG-HA-6xHis-YFP at the amino terminal position. The EMC IRES directs the expression of the IL2Rα ORF, a surface selection marker. **B,** The codon-optimized DiPRO1 ORF cloned into the pOZ vector was amplified. The primers distinguish between the exogenous (exDiPRO1) and endogenous (enDiPRO1) DiPRO1 gene. Total RNA was extracted from TE671 cells transiently transfected with exogenous DiPRO1 (pDiPRO1) and parental homologs (Ctl) 24 h post-transfection. Results represent two independent extractions of total RNA and RT-qPCR analysis performed in triplicate and normalized to GAPDH, mean ± SD. The t -test was used for statistical analyses. $^{**}P < 0.01$, $^{***}P < 0.001$ indicates difference from control. **C-E,** Myoblasts were stably transduced with a retroviral vector expressing the DiPRO1 ORF (pDiPRO1) and compared with their parental counterparts (Ctl). **C,** Heatmap revealing differential expression of a set of genes in control and pDiPRO1 samples (n=6 per condition). Upregulation of DiPRO1 gene expression (**D**) stimulates expression of PAX-family genes (**E**). **F-I,** Human myoblasts were transduced with a lentiviral vector expressing a DiPRO1-targeting shRNA (shDiPRO1) and compared to myoblasts transduced with the equivalent vector expressing a non-targeting shRNA (shCTL). The analysis was performed one week post-transduction. **F,** Cell death analysis of shDiPRO1 myoblasts. Transduced myoblasts were stained with propidium iodide (PI) and analyzed by flow cytometry. The percentage of dead cells according to DNA content was compared to control cells and the difference was estimated using the t-test, m ± SD, n=3 corresponding to three independent experiments. **G,** Heatmap revealing changes in gene expression profile related to DiPRO1 knockdown in myoblasts. (n=6-7 per condition). **H,** DiPRO1 expression was significantly inhibited by shDiPRO1 relative to shCtl. **I,** Clustered functional network of 185 upregulated (UP) and 136 downregulated (DW) genes encompassing muscle-related functions (right). Muscle GO terms are represented by nodes. Two networks were confronted. Green clusters correspond to upregulated myogenic genes, while lilac clusters correspond to downregulated genes. The kappa score = 0.4. **J,** PCA analysis of differentially expressed genes separates pDiPRO1 and control according to the first principal component and shDiPRO1 and control according to the second principal component. **A-J:** Global transcriptome analysis by microarray was implemented using total mRNA extracted from eight myoblast cell lines: with DiPRO1 knockdown (shDiPRO1) or overexpression (pDiPRO1) and corresponding controls. Three or four separate extractions were performed to collect replicates (n = 6-7 per condition). Cut-off $P<0.05$. The Limma R package was used for statistical analysis. **D, E, H:** *,**, *** indicate significant difference with the corresponding control, $P < 0.05$, 0.01, 0.001, respectively.

### Supplementary Figure S3

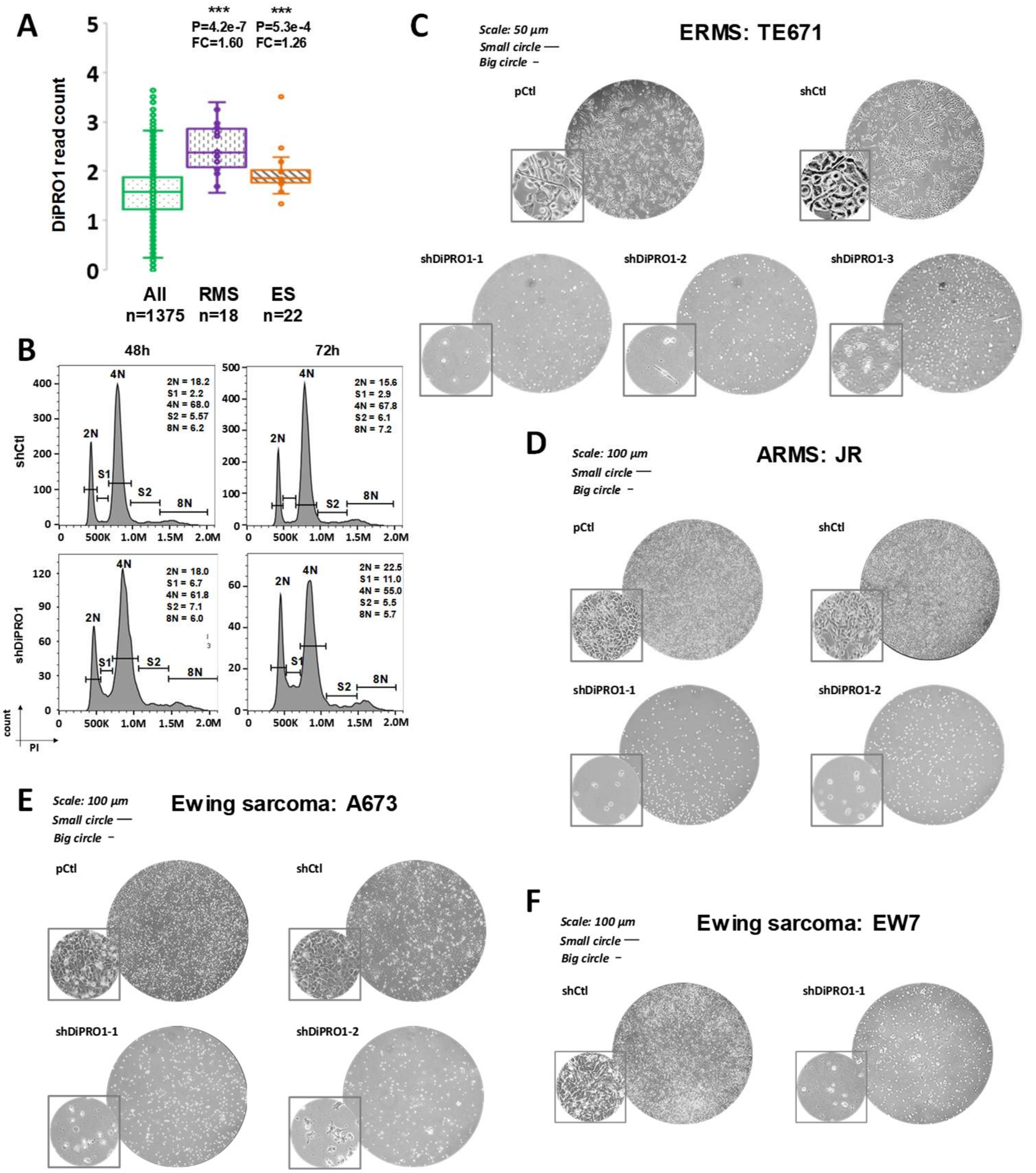

#### Supplementary Figure S3. Depletion of DiPRO1 expression affects the survival of rhabdomyosarcoma and Ewing sarcoma cells.

**A,** DiPRO1 upregulation in rhabdomyosarcoma (RMS) and Ewing's sarcoma (ES) cell lines. RNA-seq data were extracted from the DepMap portal (Broad Institute). The whisker plot shows the median (line) and quartile of ZNF555 expression by cancer type or in all cancer types (ALL). *** indicates significant difference with ALL, $P < 0.001$, t test. The number of cell lines (n) and fold change (FC) are shown in the upper part of the image. **B-F,** Stable knockdown was achieved by transduction of lentiviral vectors producing non-targeting shRNA (shCtl) or DiPRO1-targeting shRNAs (shDiPRO1-1/2/3). **B,** Cell cycle analysis of TE671 cells lacking DiPRO1 was performed 48h and 72h after transduction with vectors expressing DiPRO1-targeting shRNA. Transduced RMS cells were fixed in ethanol and stained with propidium iodide (PI). DNA content and percentage of cells in each cell cycle phase were determined by flow cytometry. The results represent two independent experiments. **C-F,** DiPRO1 knockdown induces a dramatic cell death of RMS and Ewing's sarcoma cells 5-7 days after infection. Representative images of RMS TE671 (**C**), JR (**D**) and Ewing sarcoma A673 (**E**), EW7 (**F**) cells. pCtl: untranfected parental cells, ARMS: alveolar rhabdomyosarcoma, ERMS: embryonal rhabdomyosarcoma.

### Supplementary Figure S4

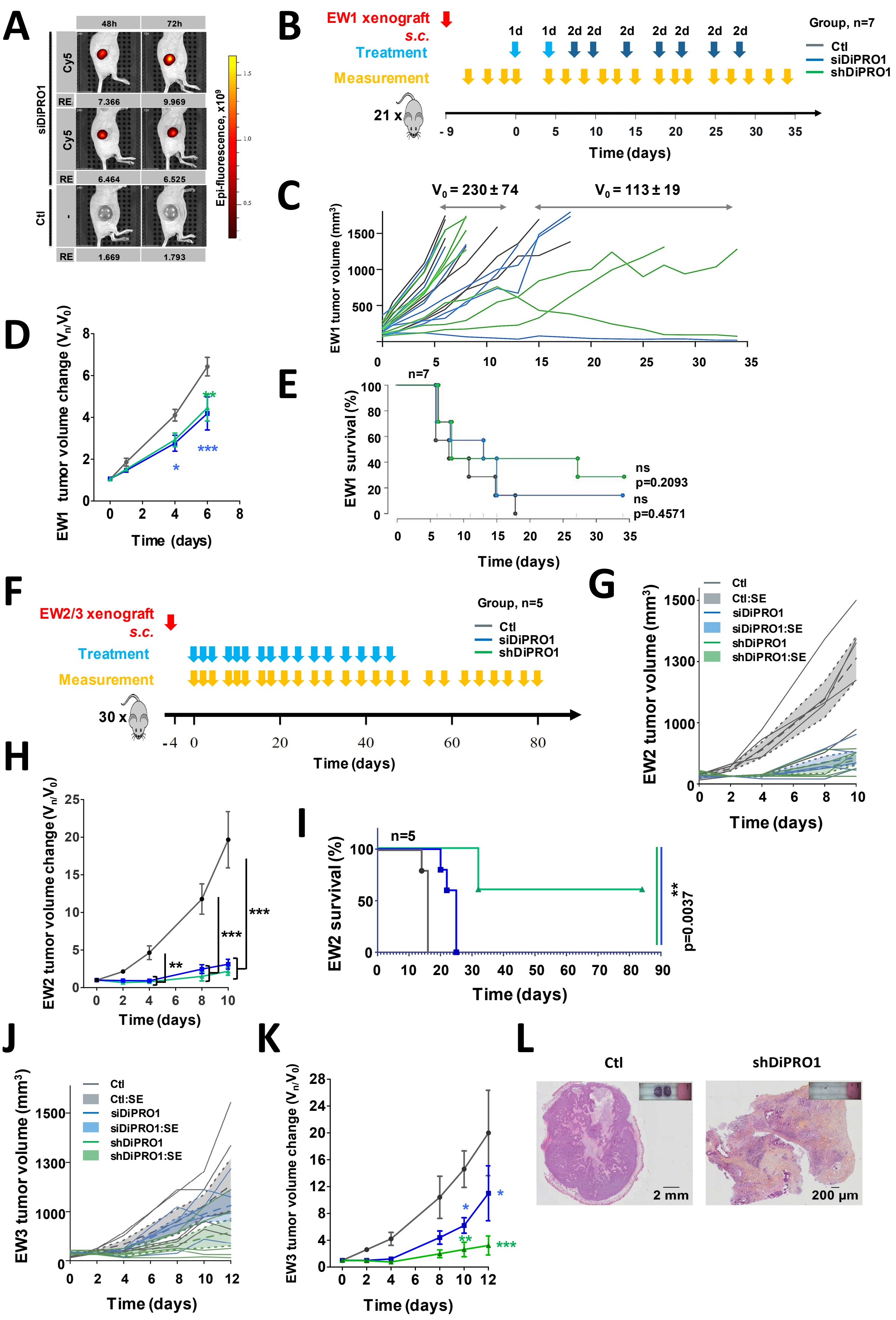

**Supplementary Fig. S4. DiPRO1 inhibitors compromise tumor growth in Ewing's sarcoma xenograft models.**

Nude mice received s.c. inoculations of A673 Ewing's sarcoma (EW) cells and were treated with siDiPRO1/jetPEI®, shDiPRO1/jetPEI®, and siCtl/jetPEI® scramble (Ctl) nanocomposites at 0.5 or 1 mg/kg/d. Three independent experiments were performed. **A,** Tumor cell uptake of siDiPRO1/jetPEI®/Cy5 nanocomposites. Specific anti-DiPRO1 siRNA complexes coupled to Cy5 were delivered at a single dose (0.5 mg/kg/) by intratumoral injection into two nude mice bearing Ewing sarcoma tumors. Internalization was monitored in live mice 24 h and 72 h post-treatment and radiant efficacy (RE) was measured. Control mice were treated with an equimolar dose of non-targeting siCtl/jetPEI® free of Cy5. **B-E,** EW1 xenograft model. **B,** Schematic showing the timing of tumor injection and treatment with DiPRO1 inhibitors. Mice were treated for 25 days at a dose of 0.5 mg/kg/inj (1d) twice weekly for the first week, and then the dose was doubled (2d) for the rest of the treatment. **C,** Growth curves of individual EW1 tumors (V, mm3). **D,** Tumor progression was compared between the treated and control groups ($V_n/V_0$, n=7). Two-way ANOVA test was used for statistical analysis. **E,** Kaplan-Meier survival curves for treated mice. Overall survival was followed for 34 days (n=7). The log-rank test (Mantel-Cox) was applied for each pairwise comparison. **F-L,** EW2 and EW3 xenograft models. **F,** Schematic showing the timing of tumor injection and treatment with DiPRO1 inhibitors. Mice were treated with siDiPRO1/jetPEI®, shDiPRO1/jetPEI®, and scrambled siRNA/jetPEI® (Ctl) nanocomposites for 80 days at a dose of 0.5 mg/kg/d (1d). The intensive three-dose-per-week regimen was used for the first two weeks, followed by a two-dose-per-week regimen until the end of treatment.. **G,** Growth curves of individual EW2 tumors (V, $mm^3$). **H,** EW2 tumor progression was compared between the treated and control groups ($V_n/V_0$, n=5). Two-way ANOVA test was used for statistical analysis. **I,** Kaplan-Meier survival curves for treated mice bearing EW2 tumor xenografts. Overall survival was followed for 84 days (n = 5). The log-rank test (Mantel-Cox) for pairwise comparison with control was applied for analysis. **J,** Growth curves of individual EW3 tumors (V, mm3). **K,** EW3 tumor progression was compared between treatment and control groups ($V_n/V_0$, n=5), means ± SEM. Two-way ANOVA test was used for statistical analysis. **L,** Representative HES staining of tumor sections from a scrambled control and a Ewing's sarcoma tumor (EW2) treated with shDiPRO1/jetPEI® shows the absence of viable tumor cells in the treated residual tumor 84 days after xenografting. **D, D, H:** All growth curves are shown as means ± SEM. * $P < 0.05$, ** $P < 0.01$, *** $P < 0.001$. P values are shown between the indicated groups and appropriate controls.

#### Supplementary Figure S5

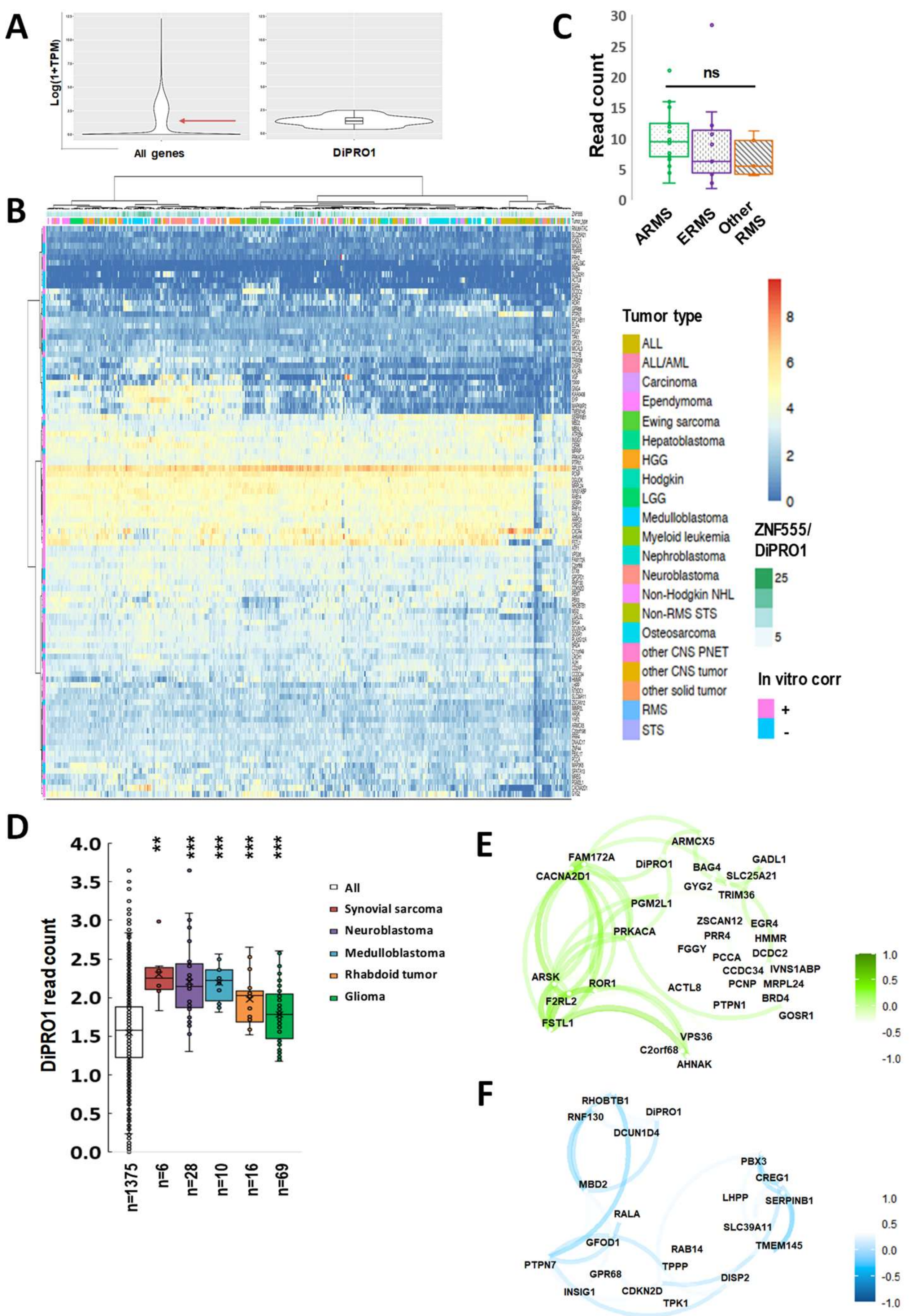

#### Supplementary Fig. S5. Clinical relevance of the DiPRO1 gene and its downstream targets in cancer.

**A,** Expression level variation of DiPRO1 among other genes in pediatric tumors. **B,** Unsupervised clustering analysis shows upregulation of the ZNF555/DiPRO1 gene in several tumor types. Genes positively (+) and negatively (-) correlated with DiPRO1 expression in in vitro DiPRO1 knockdown experiments are shown on the left side. **C,** Boxplots of DiPRO1 expression in RMS subtypes; ARMS: alveolar RMS, ERMS: embryonic RMS. Data are presented as median ± SD. Statistical analysis using the t-test showed no significant difference between subtypes. **D,** DiPRO1 overexpression in cancer cell lines. The whisker plot shows the mean (cross) and median (line) of DiPRO1 expression by cancer type or in all cancer types (ALL). The number of cell lines (n) is shown in the upper part of the image. The t-test was used for statistical analysis. **E-F,** Network plot of positive **(E)** and negative **(F)** expression correlation of DiPRO1 downstream targets across cancer cell lines. Variables that are more highly correlated appear closer together and are linked by paths (green for positive and blue for negative). The affected DEGs in RMS and Ewing's sarcoma tumor samples were analysed across cancer cell lines. **A-C:** RNA-seq data of primary tomors were processed using DEBrowser, the R package DESeq2 (TMM normalization, parametric adjustment, LRT test), and the R function network_plot (multidimensional clustering). **D-E:** RNA-seq data 1375 cancer cell lines were retrieved from the DepMap portal (Broad Institute). ns: ns: non-significant, P > 0.05, ** P < 0.01, *** P < 0.001. P values are shown between samples of the indicated cancer type and other cancer samples.

#### Supplementary Figure S6

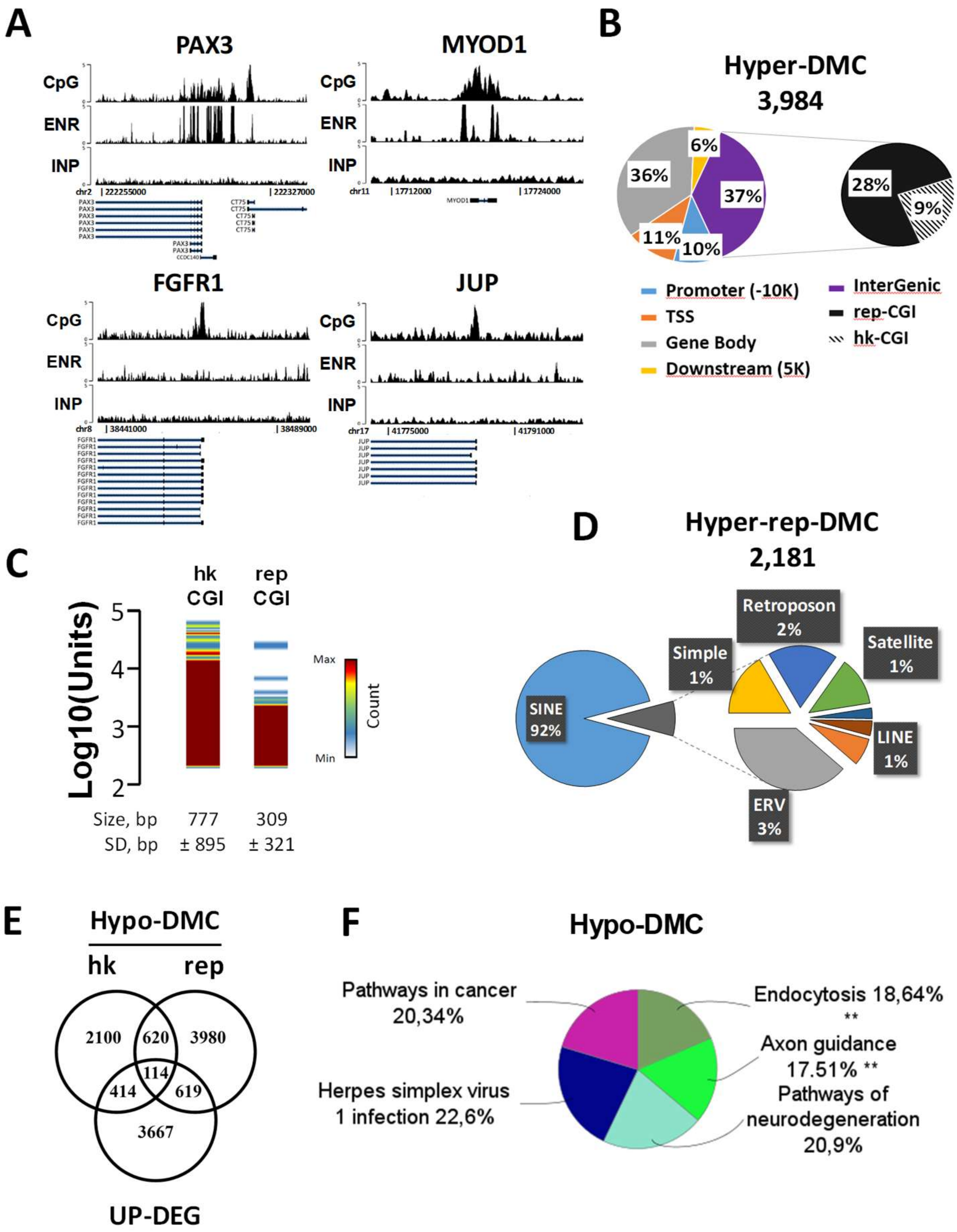

#### Supplementary Fig. S6. Methylation changes in different CGI populations related to DiPRO1 inhibition in RMS cells.

Methylation by MIRA-seq was analyzed in DNA samples from TE671 cells. Cells expressing a DiPRO1-targeting shRNA (shDiPRO1) were compared to control cells expressing a non-targeting shRNA (shCtl). Signals from methylation-enriched DNA (ENR) were normalized to the unenriched input DNA (INP). **A,** Tracking of signals on both strands corresponding to methylation signals within CGI regions of PAX3, MYOD1, FRGR1, and JUP genes in RMS control cells. Y axis represents signal intensity; X axis represents hg38 genomic coordinates. **B**, Genomic distribution of hypermethylated DMC is shown as % of DMC total number. **C**, Density plot of log-transformed size of hk- and rep-CGI in human genome. **D,** DNA repeat class distribution within DiPRO1-linked hypermethylated DMC regions. **E,** Venn diagram of overlapping genes at hypomethylated DMC associated with upregulated DEGs in RMS cells compared to control, $P<0.05$. **F,** KEGG-based functional analysis of genes associated with hypomethylated DMC. p-value of the group corrected with Bonferroni step-down, kappa score = 0.4. CGI, CpG islands according to Gardiner-Garden and Frommer criteria; DMC, differentially methylated CGI; hk-DMC, housekeeping DMC w/o repetitive elements; rep-DMC, DMC with repetitive elements.

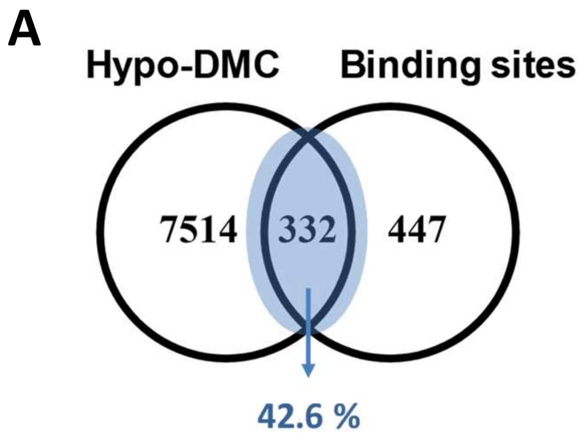


**Supplementary Fig. S7. CpG-island methylation changes in DiPRO1 binding sites**.

Venn diagrams overlapping genes associated with cis-regulatory binding sites for DiPRO1 and genes linked to differentially methylated CGIs (DMCs) in their vicinity that are hypomethylated (Hypo) or hypermethylated (Hyper) in RMS cells with DiPRO1 knockdown.

**Supplementary Table S1.** Characteristics of the patients used in the study.

| | All samples n=340 | RMS n=37 | ES n=25 |
|---|---|---|---|
| Age (years)* | 12.7±0.59 | 11.7±1.62 | 15.6±2.0 |
| Gender | | | |
| Female | 40% | 43% | 32% |
| Male | 60% | 57% | 68% |
| Treatment outcome | | | |
| Refractory | 31% | 32% | 24% |
| Relapse | 68% | 68% | 76% |

* median ± CI (95%)

**Supplementary Table S2.** Disease groups and tumor types in the study cohort.

| DISEASE GROUP | CNS TUMORS | LEUKEMIAS | LYMPHOMAS | SARCOMAS | OTHER SOLID TUMORS |
|---|---|---|---|---|---|
| TUMOR TYPE | High-grade glioma | Acute lymphoblastic leukemia | Anaplastic large-cell lymphoma | Osteosarcoma | Neuroblastoma |
| | Low-grade glioma | Acute myeloid leukemia | Other Non-Hodgkin Lymphoma | Non-rhabdmyosarcoma soft tissue sarcoma | Wilms' tumor |
| | Medulloblastoma | | | Ewing sarcoma | Hepatoblastoma |
| | Ependymoma | | | Rhabdomyosarcoma | Carcinoma |
| | Other CNS tumor | | | | Other solid tumor |
| PATIENT NUMBER | 104 | 27 | 9 | 129 | 58 |

**Supplementary Table S3.** List of primers used for transcription analysis.

| Sequence ID | Forward | Reverse |
|---|---|---|
| DiPRO1endo | TGAGCAGAATCCAGCAAAGCCCA | CCGCCTGCCCCTAGCGGTCC |
| DiPRO1exo | TCAATTTAAGGCCAGCGGGA | CCCAGCTCACGTTCCTTGTA |
| MYF5 | ATGCCATCCGCTACATCGAG | TGCCATCAGAGCAGTTGGAG |
| MYOD1 | CTCTCTGCTCCTTTGCCACA | AGTGCTCTTCGGGTTTCAGG |
| MYOG | GTGCCATCCAGTACATCGAGC | GCAGATGATCCCCTGGGTTG |
| MYH1 | GAGTCCATGCAGAGCACACT | GCTGTTCCTTCAGGTCCTCC |
| GAPDH | CCAGGTGGTCTCCTCTGACT | AAGTGGTCGTTGAGGGCAAT |
